# A critical assessment of aptamer and CRISPR-Cas12a-based biosensors for small molecule detection

**DOI:** 10.1101/2025.10.17.683058

**Authors:** Oliver F. Brandenberg, Elisabeth M.-L. Janssen, Olga T. Schubert

## Abstract

Analyte detection through aptamer-induced signal generation by CRISPR-Cas enzymes has rapidly emerged as a popular biosensing approach. Here, we investigated the implementability and analytical performance of this setup for the detection of diverse small molecule analytes. We selected nine aptamers from the literature targeting seven analytes and tested a commonly used assay design whereby analyte binding by the aptamer liberates a short complementary DNA strand, which in turn activates Cas12a to generate a fluorescence signal. After extensive optimization, the assay functioned for only two of the seven analytes, and several previously reported results could not be reproduced. While Cas12a fluorescence detection was robust, the low success rate is likely due to aptamers not functioning reliably, underscoring the need for careful aptamer validation. Overall, our study provides a critical assessment of aptamer-Cas12a assay performances and discusses potential strengths, limitations, and pitfalls of this biosensing strategy.

## Introduction

Rapid detection and quantification of diverse analytes is a critical need across different scientific disciplines. In medicine, point-of-care testing to detect disease agents or to measure biomarkers is an important component of modern healthcare.^1^ In agriculture and the food industry, detection of toxins or unwanted components in agricultural products or processed consumables is essential for consumer safety.^2^ And in the environmental sciences, detection of pollutants in soil, water, or air is critical to enforce environmental protection standards and to monitor human exposure limits.^3^

Liquid or gas chromatography coupled to mass spectrometry (LC-MS or GC-MS, respectively) represent the state of the art for accurate detection and quantification of analytes ranging from small molecules to proteins. While those methods allow simultaneous analysis of thousands of compounds, they are often time-consuming and require trained personnel as well as specialized, expensive equipment, which limits their availability, utility for field testing, and sample throughput.^4,5^

Alternative analytical techniques that can overcome some of the limitations of MS-based methods include biosensors. A biosensor typically consists of two components: The first component is a bioreceptor, that is, a biomolecule capable of selectively binding the analyte of interest; to date, the most widely used bioreceptors are antibodies. In a second step, a transducer component translates the bioreceptor-analyte interaction into a measurable signal; common signal transducers include optic (colorimetric, fluorescent, chemiluminescent, surface plasmon resonance) or electrochemical (voltammetric, amperometric, potentiometric) devices.^6,7^

In the early 1990s a new class of bioreceptors emerged, called aptamers. Aptamers are short (typically 30 to 100 nucleotides long) single-stranded DNA or RNA molecules capable of binding an analyte with high affinity and specificity. Aptamers for a given analyte are selected from a large starting library by an *in vitro* procedure called systematic evolution of ligands by exponential enrichment (SELEX).^8,9^ Aptamers possess several favorable properties, including no need for animal immunizations, low-cost synthesis, and high stability (particularly when lyophilized), rendering them excellently suited for biosensor applications, especially in field settings.^10,11^ Hence, in the last three decades, aptamers targeting a wide range of analytes have been developed and subsequently employed to construct diverse biosensors, commonly referred to as aptasensors.^12,13^

A recent addition to aptasensor-based biosensors is the use of CRISPR-Cas enzymes as the transducer component for signal generation.^14,15^ In this setup, a short DNA strand (hereafter referred to as activator DNA) is annealed to the aptamer by complementary base pairing to yield an aptamer-activator DNA duplex. Upon binding of the analyte to the aptamer, the activator DNA is displaced and remains free in solution, where it can be detected. This principle of target-induced DNA strand displacement has already been utilized for aptasensor development over the past 20 years.^16^ The novelty of the CRISPR-Cas aptasensor format is that the free activator DNA activates a CRISPR-Cas enzyme by binding to its suitably designed, complementary CRISPR-RNA (crRNA). The most frequently used CRISPR-Cas enzyme in this setup is Cas12a (Cpf1); once activated, this enzyme possesses indiscriminate single-stranded DNA (ssDNA) cleavage activity.^17^ This cleavage activity is exploited by supplying a short ssDNA reporter labelled with a fluorophore and a quencher, and cleavage of this reporter by activated Cas12a generates a fluorescence signal. Thus, by coupling analyte binding by the aptamer to activator DNA release and subsequent Cas12a activation, this setup allows the detection of low concentrations of analyte through an enzyme-amplified fluorescence signal (Figure 1).

**Figure 1.**
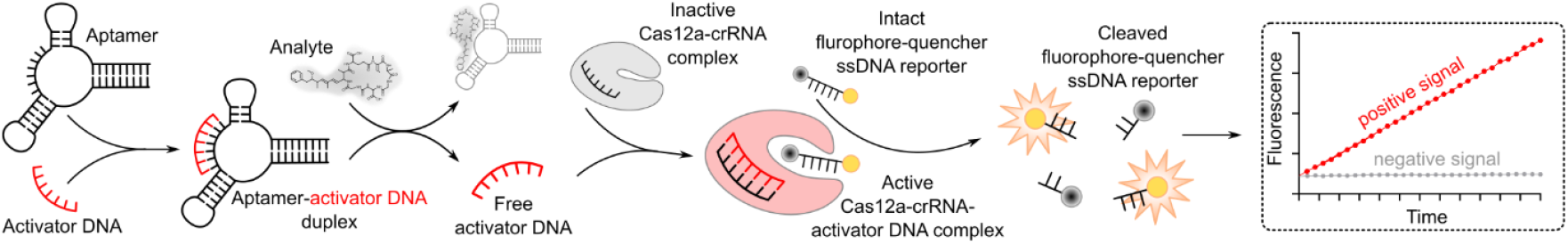
Overview of the Cas12a-aptasensor assay. An aptamer is annealed with a 20 nt-long activator DNA to yield an aptamer-activator DNA duplex. Upon analyte binding by the aptamer, the activator DNA is released. The free activator DNA can then bind to the crRNA of an inactive Cas12a enzyme, thereby activating it. Activated Cas12a will cleave a short ssDNA labelled with a fluorophore and a quencher; cleavage of this reporter will generate a fluorescence signal that can be followed either kinetically (shown) or as an endpoint assay.

Since the first publication of this aptamer-Cas12a biosensor approach in 2019,^18,19^ more than 150 studies have reported modified versions of this setup to construct biosensors for diverse analytes (Table S1). Often, these assays can be performed in 96-well plate format with a turnaround time of 60 to 120 minutes, allowing high-throughput sample analysis. In addition, due to the Cas12a enzyme-amplified signal generation, analyte limits of detection and quantification (LOD and LOQ, respectively) in the low nanomolar range have frequently been reported. It is often argued that this strategy is universally applicable, provided that suitable aptamers for the analyte of interest are available (Table S1 and references therein).

Indeed, one major prerequisite for this aptasensor approach is the availability of an aptamer that, upon binding its cognate analyte, efficiently releases the activator DNA. Typically, this requires structure-switching aptamers, *i*.*e*., aptamers undergoing a conformational change upon target binding. Structure-switching may be an adventitious property of an aptamer; alternatively, such aptamers can be specifically obtained through a modified SELEX protocol called Capture-SELEX.^20,21^ Here, structure-switching aptamers are selected through analyte-induced dissociation of the aptamer from a short ssDNA (10-15 nucleotides long) complementary to the aptamer. Thus, Capture-SELEX aptamers are, in principle, well suited for the CRISPR-Cas aptasensor approach, as the sequence of the capture strand can be directly adapted as activator DNA. Over fifty Capture-SELEX aptamers against diverse small molecule analytes have been reported in recent years, providing a broad resource to develop CRISPR-Cas aptasensors.^22^

Considering this large pool of published structure-switching aptamers and the touted benefits of the CRISPR-Cas aptasensor setup, we set out for a proof-of-concept study. In our daily work as environmental analytical chemists we measure many different water contaminants by LC-MS.^23–25^ Thus, we were curious to evaluate whether the CRISPR-Cas aptasensor technology could provide a potential alternative for facile and sensitive small-molecule detection. Specifically, we had the following questions: Is it possible to select structure-switching aptamers for different analytes from the literature and use them in the advertised plug-and-play manner for the CRISPR-Cas aptasensor setup? Can we reproduce assay performance of already published CRISPR-Cas aptasensors? And what are potential pitfalls and limitations when implementing this aptasensor setup?

We systematically addressed these questions in several steps. First, we selected a test set of nine aptamers targeting seven small molecule analytes; one of those had previously been published as a CRISPR-Cas aptasensor. We then systematically optimized the Cas12a fluorescence read-out, determined the quantification limits of the activator DNAs, investigated the stability of the aptamer-activator DNA duplexes, and tested two different formats of the CRISPR-Cas aptasensor assay. We found that, in our hands, the assay worked for only two out of the seven analytes we investigated. Regarding this low success rate, we provide a critical analysis of this experimental setup as well as a general discussion of the potential limitations for implementing the CRISPR-Cas aptasensor strategy.

## Results

### Selection of aptamer-analyte combinations and design of activator DNAs

To test and implement CRISPR-Cas aptasensors, we selected nine published aptamers targeting seven analytes (Figure 2):

1. Aptamer AN6 obtained by classical SELEX and targeting the cyanobacterial toxin microcystin-LR (MC-LR), with a reported dissociation constant (K_d_) of 50 nM.^26^ This aptamer had previously been used in CRISPR-Cas aptasensor setups with reported MC-LR limits of detection (LOD) of 3 pg/L (3 fM)^27^ and 4.5 ng/L (4.5 pM).^28^
2. Aptamer R12.45 obtained by Capture-SELEX and targeting the herbicide atrazine, with a reported K_d_ of 3.7 nM.^29^
3. Aptamers FipA8 and FipA6B obtained by Capture-SELEX and targeting the insecticide fipronil, with a reported K_d_ of 2 and 0.9 nM, respectively.^30^
4. Aptamer PQ4 obtained by Capture-SELEX and targeting the antimalarial drug piperaquine, with a reported K_d_ of 2.4 nM.^31^
5. Aptamers Q2 and Q8 obtained by Capture-SELEX and targeting the antibiotic ofloxacin, with a reported K_d_ of 0.1 and 0.2 nM, respectively.^32^
6. Aptamer LxCsh, an RNA aptamer obtained by Capture-SELEX and targeting the antibiotic levofloxacin (the S-isomer of ofloxacin), with a reported K_d_ of 6 μM.^33^
7. Aptamer MN19 obtained by classical SELEX and targeting the antimalarial drug quinine (though originally selected against cocaine), with a reported K_d_ of 700 nM.^34^ A variant of this aptamer (MN4) had previously been used for CRISPR-Cas aptasensor assays for cocaine.^35,36^ This very well studied aptamer is a canonical example of structure-switching aptamers and was included here as a benchmark.

**Figure 2.**
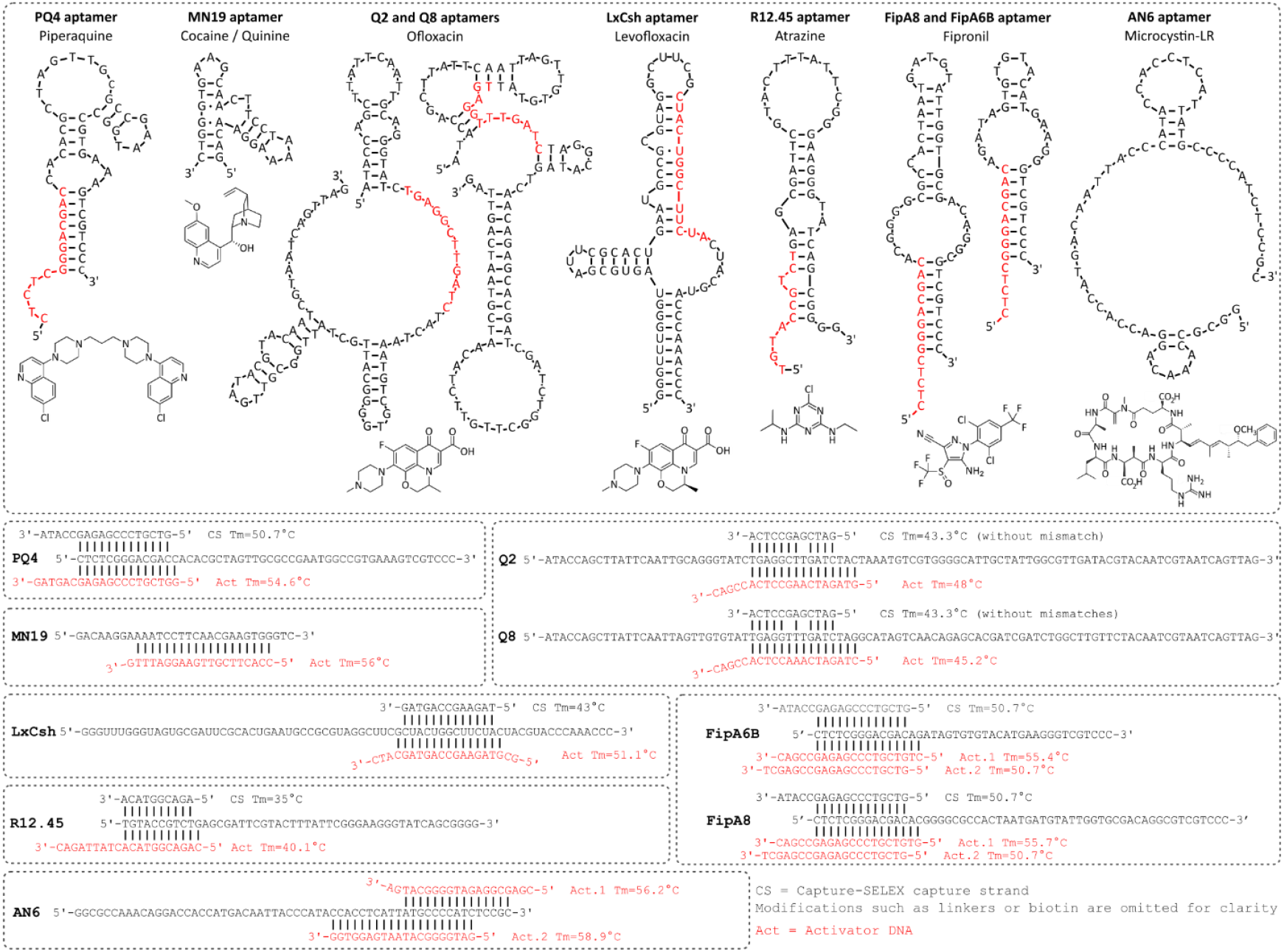
Selected aptamers and corresponding capture strands and activator DNAs. Top: Nucleotide sequences and secondary structures of the aptamers used in this study together with the molecular structures of their corresponding analytes. Aptamer secondary structures were adapted from the original publications except for aptamer AN6, which was predicted using mfold.^37^ For aptamers obtained by Capture SELEX, the sequence complementary to the capture strand is shown in red. Bottom: Linear depiction of aptamer sequences together with their Capture-SELEX capture strands (if applicable) and the activator DNAs; the melting temperature of the capture strand (CS Tm) and activator DNA (Act Tm) is also indicated. Activator DNAs were designed *de novo* except for MN19 and AN6, which were obtained from the literature.^27,28,35^

We designed matching activator DNAs and crRNAs for all selected aptamers (Table S2). For aptamers obtained through Capture-SELEX we designed the activator DNA based on the ssDNA capture strand used in the Capture-SELEX campaign. In all those cases, the length of the capture strand was between 10 and 15 nt. As the activator DNA length for optimal Cas12a activation is 20 nt, we added random non-annealing nucleotides to achieve this length for all activator DNAs. For aptamers previously used in CRISPR-Cas aptasensor assays, we designed the activator DNAs exactly as described in those studies.^27,28,35^ The position and sequence of the capture DNA strands (if applicable) and activator DNAs is shown in Figure 2, and the matching crRNAs are displayed in Figure S1.

### Optimization of the Cas12a fluorescence assay

The reliable and sensitive detection of free activator DNA through a Cas12a-generated fluorescence signal is a prerequisite for the CRISPR-Cas aptasensor workflow. Thus, we first worked towards optimizing the Cas12a-based fluorescence assay with respect to buffer composition, enzyme concentration, and fluorophore-quencher ssDNA reporter length. It was reported that the activity of Cas12a can be improved by substituting the buffer supplied by the manufacturer (buffer NEB2.1) with different buffers or buffer additives.^38^ This includes substitution of chlorine-based salts with acetate or sulfate salts, addition of the reductant dithiothreitol (DTT), addition of bovine serum albumin (BSA; already present in NEB2.1), and substitution of the divalent cation Mg^2+^ with Mn^2+^.^38 39 40^ We thus tested several standard buffers (NEB2.1, PBS, TE; see Table S3 for all buffer compositions) and various alternative buffers for their effect on Cas12a activity (Figure S2A, B). Buffer CM, containing 10 mM Mn^2+^, resulted in the highest Cas12a fluorescence generation rate (4.7-fold higher than with buffer NEB2.1). However, we also repeatedly observed a brown precipitate forming with this buffer, presumably caused by the oxidation of Mn^2+^, which we deemed problematic for reproducible assay performance. The second-best buffer, called CXB herein, was derived from the CEXTRAR buffer described by Habimana *et al*.^38^ and contained 50 mM potassium acetate, 20 mM Tris-acetate pH 8.0, 20 mM MgSO_4_, 10 mM DTT, and 0.1 mg/mL BSA. This buffer resulted in a Cas12a fluorescence generation rate 3.6-fold higher compared to buffer NEB2.1 (Figure S2B). Unless otherwise noted, this buffer was employed for all subsequent CRISPR-Cas aptasensor assays.

We next tested different Cas12a enzyme concentrations (10 nM, 20 nM, 40 nM) with equimolar amounts of crRNA. A 10 nM concentration of Cas12a-crRNA complex provided the highest cleavage activity while also representing the most economical setup and was chosen for all subsequent assays (Figure S2C).

Most previous Cas12a-aptasensor studies (Table S1) used a 5 nt-long fluorophore-quencher ssDNA reporter to detect Cas12a activity. However, it was reported that a longer ssDNA reporter (up to 13 nt) may increase Cas12a cleavage rates.^38^ We thus tested a 12 nt fluorophore-quencher ssDNA reporter alongside the 5 nt reporter. While the Cas12a cleavage rate of the 12 nt reporter was indeed slightly higher, the background level also increased, presumably due to less efficient quenching (Figure S2 D, E). We thus continued our work with the 5 nt ssDNA reporter. We set the reporter concentration at 500 nM (a 50-fold molar excess over the maximum theoretical concentration of active Cas12a-crRNA complexes) to avoid substrate limitation and to enable us to measure stable Cas12a enzyme kinetics.^40^

We set the Cas12a assay temperature at 25°C. Higher reaction temperatures (up to 45°C) were reported to result in higher Cas12a activity.^41^ However, higher temperatures may interfere with the aptasensor assay by inducing activator DNA dissociation from the aptamer even in the absence of analyte. A reaction temperature of 25°C still enables high Cas12a activity while being well below the calculated melting points of our aptamer-activator DNA pairs (Figure 2), ensuring that any free activator DNA only originates from aptamer-analyte interactions.

We further tested whether activator DNAs shorter than the canonical 20 nt would activate Cas12a, as this would provide more flexibility in activator DNA design. Thus, we designed and tested a set of shorter activator DNAs for atrazine, which revealed that only the full-length 20 nt sequence efficiently activated Cas12a (Figure S4), leading us to use 20 nt activators in all subsequent assays.

### Determination of limits of detection and quantification for activator DNAs in the Cas12a assay

Having established ideal conditions for the Cas12a fluorescence assay, we next determined LODs and LOQs for all our activator DNAs. To this end, we performed two-fold dilution series of each activator DNA from 4 nM to 1.95 pM and measured the resulting fluorescence generation by Cas12a. We used the slopes of fluorescence increase over time (instead of end points) to calculate LODs and LOQs by standard linear regression procedures. Exemplary data for the atrazine activator DNA are shown in Figure 3A and B; fluorescence data and linear regression analyses of all other activators are shown in Figure S3. For all activators, we obtained LODs and LOQs in the low picomolar range (Figure 3C). This bodes well for the potential LODs and LOQs of the analytes in the Cas12a-aptasensor assay: In the simplest case, binding of a target molecule to an aptamer–activator DNA duplex can be assumed to follow a 1:1 stoichiometry, such that each target molecule releases one activator DNA. Accordingly, the obtained activator DNA LODs would directly reflect the LODs for the actual target analytes.

**Figure 3.**
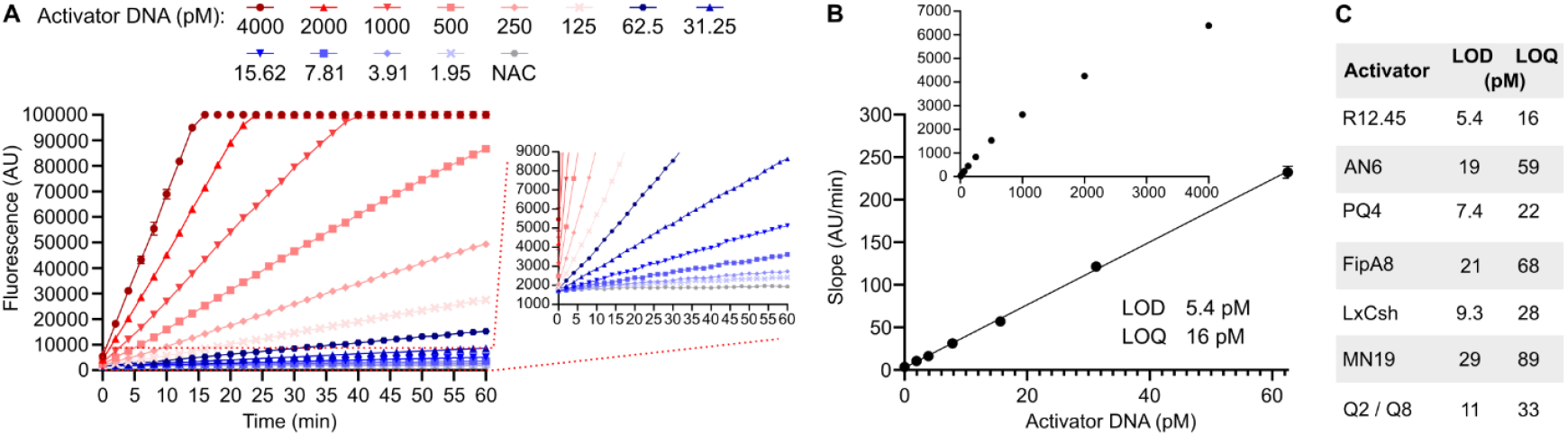
Activator DNA LOD and LOQ in the Cas12a fluorescence assay. (A) Dilution series of the atrazine R12.45 activator DNA from 4 nM to 1.95 pM and the resulting fluorescence signals generated in the Cas12a assay. Data are from two independent experiments, each performed in duplicates. Inset: Linear increases in fluorescence over time could be observed even for low picomolar activator DNA concentrations. NAC, no activator DNA control. (B) Slope of fluorescence increase over time derived from the data shown in (A). Linear regression was performed on the linear part of the data (typically below 250 pM activator DNA) to derive LOD and LOQ values. (C) The obtained LOD and LOQ for all activator DNAs used in this study.

### Quantification of aptamer-activator DNA complex formation and stability

We next investigated duplex formation between all aptamers and their respective activator DNAs. In the Cas12a aptasensor assay, any free activator DNA will elicit a positive signal and thus should, in the absence of analyte, be fully sequestered by its matching aptamer. The amount of activator DNA that remains free and non-bound by aptamer depends on the molar ratio between aptamer and activator DNA and thermodynamic factors (such as the melting temperature of the duplex and structural properties of the aptamer), which differ for each aptamer-activator DNA pair (Figure 2). Thus, to investigate the optimal ratio of aptamer to activator DNA required to achieve activator DNA sequestration, we combined a fixed amount of activator DNA (10 nM) with different amounts of aptamer (molar aptamer-activator DNA ratios of 0:1, 1:1, 2:1, 4:1, 8:1), annealed them, and tested the solution for presence of free activator DNA using the Cas12a fluorescence assay (Figure 4A). Exemplary data for the ofloxacin Q2 aptamer-activator pair are shown in Figure 4B, indicating that an excess of aptamer (8:1 aptamer-activator DNA ratio) is necessary to ensure that almost no free activator DNA remains in solution. This is in agreement with the literature and was true for all our aptamer-activator pairs (Figure S5).^30,31^

**Figure 4.**
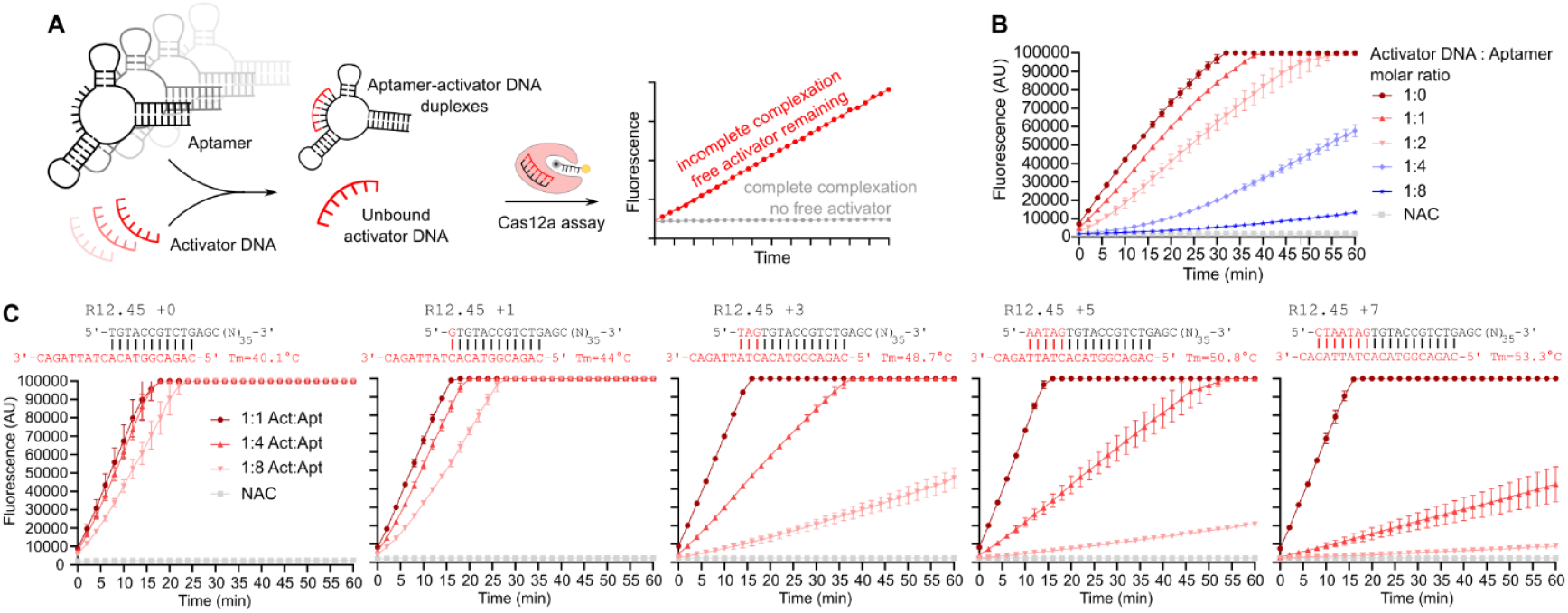
Aptamer-activator DNA duplex formation. (A) Scheme of the assay setup: Different molar ratios of aptamer and activator DNA were annealed, and the solution was tested for the presence of free activator DNA with the Cas12a fluorescence assay. (B) Exemplary data for the ofloxacine Q2 aptamer-activator DNA pair. At a molar ratio of 8:1 aptamer:activator DNA, only low amounts of activator DNA remain in solution. (C) Relationship between aptamer-activator DNA duplex melting temperature (T_m_) and duplex formation efficiency, demonstrated with the atrazine R12.45 aptamer and activator DNA. As the complementary region is increased from 11 nt to 18 nt through a stepwise elongation of the aptamer, duplex formation becomes increasingly more favorable

We also used this experimental setup to investigate the relationship between calculated aptamer-activator DNA duplex melting temperature (T_m_) and sequestration efficiency. The activator DNAs reported in CRISPR-Cas aptasensors (Table S1), as well as the capture DNA strands used in Capture-SELEX, typically contain 10-15 bases complementary to the aptamer, resulting in T_m_’s between 40°C and 60°C (Figure 2). For both Capture-SELEX and CRISPR-Cas aptasensors, it is crucial to strike a delicate balance: the aptamer-activator DNA duplex has to be sufficiently stable in the absence of the analyte but efficiently dissociate in its presence.

We tested the relationship between predicted T_m_ and aptamer-activator DNA duplex formation using the atrazine R12.45 aptamer. The capture strand used in the atrazine Capture-SELEX campaign anneals at the 5’ end of the aptamer and has a length of 10 nt and a predicted T_m_ of 35°C.^29^ We designed our activator DNA accordingly, with an additional base pair at the activator DNA 5’ end and a 9 nt non-annealing overhang at the 3’ end, for a length of 20 nt and a predicted T_m_ of 40.1°C (Figure 2, Figure 4C). However, this activator DNA did not form a stable complex with the R12.45 aptamer, as free activator DNA remained in solution even in presence of an 8-fold excess of aptamer (Figure 4C). Thus, we performed a stepwise elongation of the R12.45 aptamer at the 5’ end to anneal with the 3’ overhang of the activator DNA, creating a series of aptamer-activator DNA pairs with increasing T_m_. This revealed that an annealing length of 15 nt with a predicted T_m_ of 50.8°C was required to minimize free activator DNA in solution at an 8:1 aptamer:activator DNA ratio (Figure 4C). Similar but less pronounced effects were observed for other aptamer-activator pairs (*e*.*g*., the LxCsh, FipA8 and FipA6B aptamers),^30,33^ whereby the activator DNAs designed based on the original Capture-SELEX ssDNA reporter were not very efficient in forming a complex with the matching aptamers (Figure S5). In such cases, we designed activator DNAs based both on the original Capture-SELEX strands and longer versions with slightly elevated T_m_, and tested both activator DNAs in subsequent CRISPR-Cas aptasensor assays.

### Effect of target substances on the Cas12a fluorescence assay

In most CRISPR-Cas aptasensor assay formats, aptamer-activator DNA complexes are first incubated with analyte and the solution is then analyzed with the Cas12a fluorescence assay. Hence, the analyte is present during the Cas12a reaction and may potentially interfere with fluorescence generation. Therefore, we tested whether the seven analytes tested here had an influence on Cas12a activity. A high analyte concentration (50 μM for MC-LR, 100 μM for all other analytes) was added to a standard Cas12a assay and the effect on fluorescence generation was monitored (Figure S6). Most analytes had only minor effects on Cas12a activity, while two analytes had more pronounced effects: quinine reduced the rate of fluorescence generation, while piperaquine increased the rate of fluorescence generation. All other substances, including DMSO and methanol solvent controls, had only minor effects on Cas12a activity. For quinine and piperaquine, a dilution series revealed that in both cases, concentrations of 25 μM and lower had only negligible effects (Figure S6) and should thus not influence the results of the Cas12a read-out.

### Evaluation of the one-pot CRISPR-Cas aptasensor assay

After optimizing the CRISPR-Cas12a fluorescence assay and aptamer-activator DNA duplex formation, we proceeded to test the simplest implementation of the CRISPR-Cas aptasensor assay. Here, the aptamer-activator DNA duplex is incubated with analyte and the sample is then directly analyzed with the Cas12a fluorescence assay; we refer to this assay as one-pot setup as, in principle, the assay can be performed in a single tube (Figure 1).

First, we annealed aptamer and activator DNAs as previously established, using 25 nM activator DNA and an 8-fold molar excess (200 nM) of aptamer. We then added analyte or solvent controls and incubated the samples for 30 min at different temperatures, ranging from 25 to 40°C. We included these different incubation temperatures to increase the chances of finding a “sweet spot” where the aptamer-activator DNA duplex remains stable in the absence of analyte, but sufficiently labile to dissociate and release the activator DNA in its presence. Following the incubation, the samples were analyzed by the Cas12a fluorescence assay.

A stronger fluorescence signal is expected in samples containing the analyte compared to control. However, we observed such fluorescence differences only for two out of seven aptamer-analyte pairs: PQ4-piperaquine and MN19-quinine (Figure 5 A, B).

**Figure 5.**
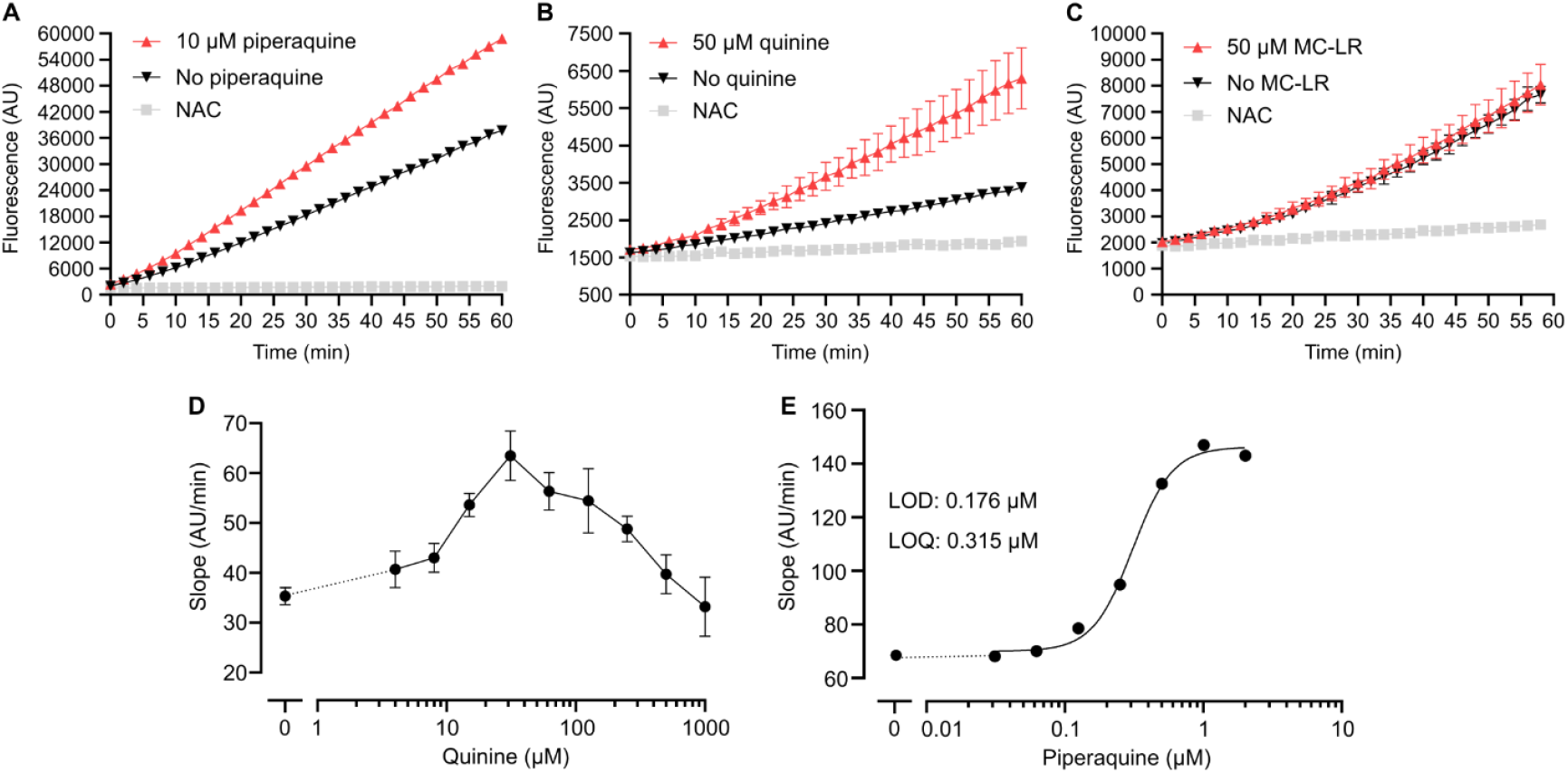
One-pot CRISPR-Cas aptasensor assay evaluation. (A) Data for the CRISPR-Cas aptasensor assay with piperaquine, showing a pronounced difference in fluorescence generation between piperaquine and control samples, indicating activator DNA release from the aptamer by piperaquine. (B) Data for the Cas12a-aptasensor assay with quinine. (C) Example of negative assay data for the analyte MC-LR. No difference in fluorescence generation between MC-LR and control samples is observable, indicating that no activator DNA is released from the aptamer in presence of MC-LR. (D) CRISPR-Cas aptasensor assay data with a quinine dilution series. Shown are the processed data (slope of fluorescence increase over time versus quinine concentration), showing a bell-shaped curve due to the inhibition of the Cas12a assay by high quinine concentrations. (E) CRISPR-Cas aptasensor data with a piperaquine dilution series, showing a sigmoidal dose-response relationship between fluorescence increase over time and piperaquine concentrations; the curve fit as well as LOD and LOQ are shown.

To achieve a signal for all remaining aptamer-analyte pairs, we engaged in extensive assay optimization. This included testing a variety of different assay buffers, increasing the incubation temperature up to 50°C, using different amounts of aptamer-activator DNA duplexes, as well as increasing the incubation time from 30 min up to 120 min. This often resulted in higher signals for both samples with and without analyte, but with no resolution between the two (Figure 5C). We therefore concluded that, in our hands, the simple one-pot CRISPR-Cas aptasensor assay worked for only two out of seven analytes (two out of nine aptamers) tested.

For quinine and piperaquine, the two functioning aptamer-analyte pairs, we characterized assay performance by testing serial dilutions of these analytes. For quinine, this was complicated by the fact that quinine concentrations above 25 µM suppress fluorescence signal generation in the Cas12a assay as discussed above (Figure S6). This resulted in a dose-response relationship reminiscent of a bell curve rather than the expected sigmoidal curve (Figure 5D). While this assay is thus not analytically useful, it demonstrates that the MN19-quinine one-pot CRISPR-Cas aptasensor, in principle, functions. For the piperaquine one-pot CRISPR-Cas aptasensor assay, we obtained the expected sigmoidal dose-response curve for concentrations from 2 to 0.03125 µM piperaquine (Figure 5E). LOD and LOQ were determined as 0.176 and 0.315 µM, respectively.

### Evaluation of the two-pot magnetic bead CRISPR-Cas aptasensor assay

In a second line of assay establishment, we tested a two-pot setup, where aptamer-activator DNA duplex incubation with the analyte and the Cas12a read-out are separated. By using a biotinylated aptamer, the aptamer-activator DNA duplex is immobilized on streptavidin-functionalized magnetic beads (Figure 6A). After incubating the duplex-loaded beads with analyte, the beads, including target-bound aptamers and remaining aptamer-activator DNA duplexes, are removed and the remaining solution, expected to contain the detached activator DNA, is transferred to the Cas12a fluorescence assay. The advantage of this setup is that no additional activator DNA can be released during the Cas12a read-out, potentially reducing background noise. Furthermore, aptamers can be annealed with activator DNA in a 1:1 molar ratio, as any unbound activator DNA can be removed during bead preparation. In contrast to the one-pot assay, which requires a large molar excess of aptamer to ensure the absence of free activator DNA, this ensures that mostly functional aptamer-activator DNA duplexes are present for analyte incubation.

**Figure 6.**
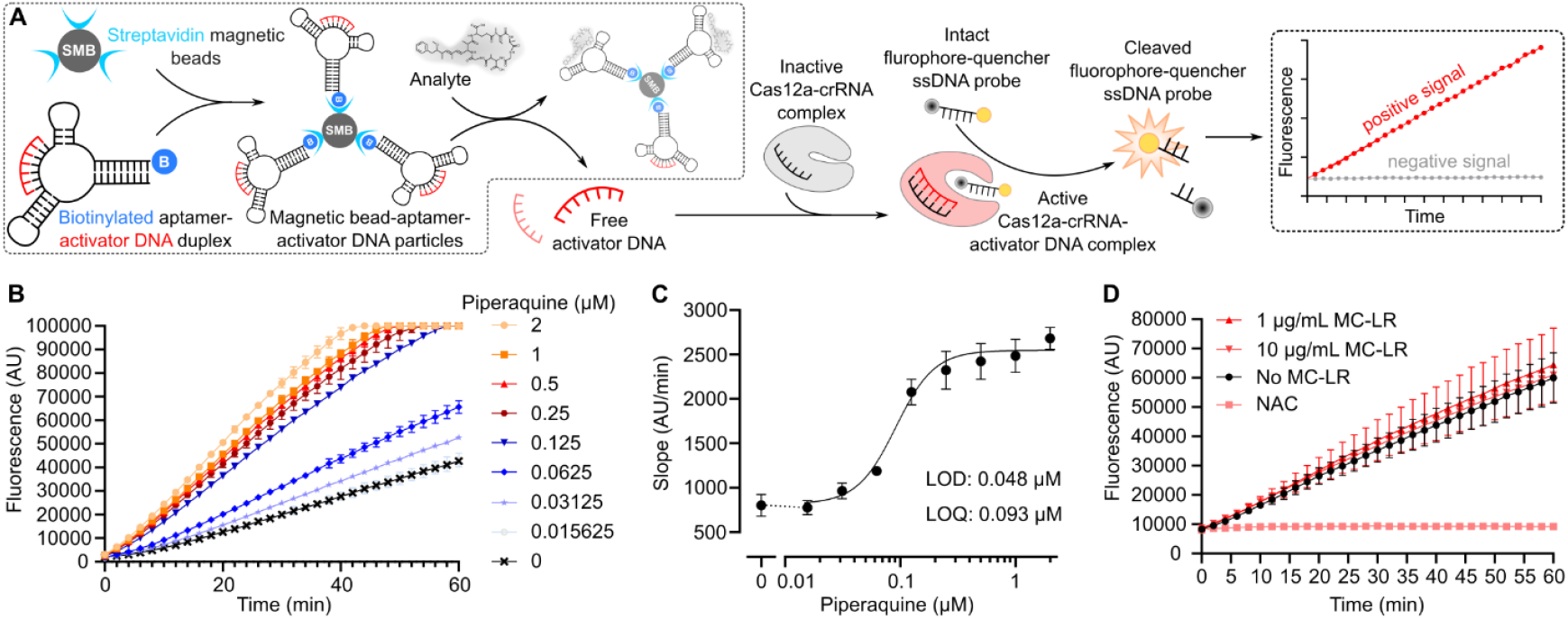
Two-pot CRISPR-Cas aptasensor assay evaluation. (A) Assay overview showing how aptamer-activator DNA duplexes are coupled to magnetic beads, which allows for spatial and temporal separation of aptamer-activator DNA duplex incubation with analyte and the subsequent Cas12a fluorescence assay. (B) Two-pot CRISPR-Cas aptasensor data for a piperaquine dilution series showing analyte concentration-dependent fluorescence generation. Data from one exemplary assay performed in duplicates are shown. (C) The slope of fluorescence changes over time versus piperaquine concentration, derived from two independent experiments as shown in (B), revealing a sigmoidal dose-response curve. The curve fit and LOD and LOQ are shown. (D) Exemplary two-pot CRISPR-Cas aptasensor data for MC-LR, where we did not observe changes in fluorescence generation for samples with or without MC-LR present.

We tested this two-pot setup with piperaquine, which functioned well in the one-pot CRISPR-Cas aptasensor setup, and with microcystin-LR, an analyte previously reported to be detectable using this exact two-pot setup.^27^ We used aptamer-activator DNA duplex-modified magnetic beads corresponding to a theoretical duplex concentration of 100 nM per assay. We then subjected the functionalized beads to samples containing analyte or solvent controls and incubated them for 30 to 120 min at 25ºC and 38ºC for piperaquine and MC-LR, respectively. Subsequently, the beads were removed magnetically, and the supernatant was subjected to the Cas12a assay.

For piperaquine, the two-pot CRISPR-Cas assay functioned as expected, and we obtained dose-dependent fluorescence signals (Figure 6B) and a sigmoidal dose-response curve (Figure 6C), with LOD and LOQ determined at 0.048 and 0.093 µM, respectively.

Although this aptasensor assay format was previously published for MC-LR with an excellent reported analytical performance and a MC-LR LOD as low as 3 fM,^27^ in our hands, these data were not reproducible. Specifically, we were unable to detect a difference in fluorescence signals between samples with or without MC-LR. Notably, we performed numerous attempts at assay optimization, including testing a large array of buffers and assay conditions and testing of aptamer, activator DNA, and MC-LR batches ordered from different suppliers. However, depending on the chosen conditions, we either received very low signals for both MC-LR-containing and control samples, or a high background fluorescence for both samples (Figure 6D).

### Comparison of aptasensor assay performances

The CRISPR-Cas aptasensor assay is based on analyte-induced activator DNA dissociation from the aptamer; indeed, this property can also be (and often has been) utilized for a simpler assay setup. In cases where the activator DNA (or original Capture-SELEX strand) anneal at one of the termini of the aptamer, the aptamer can be labelled with a fluorophore, while the complementary strand is labelled with a quencher. When the two modified DNA molecules form a duplex, fluorescence is quenched by the proximity of the fluorophore and quencher; upon analyte binding, the quencher strand is released, resulting in a fluorescence signal (Figure 7A).^42^ Whereas the CRISPR-Cas aptasensor detects free activator DNA via an enzyme-amplified kinetic fluorescence signal, this simpler fluorophore-quencher displacement assay (hereafter referred to as F-Q assay) generates a non-amplified endpoint fluorescence signal.

**Figure 7.**
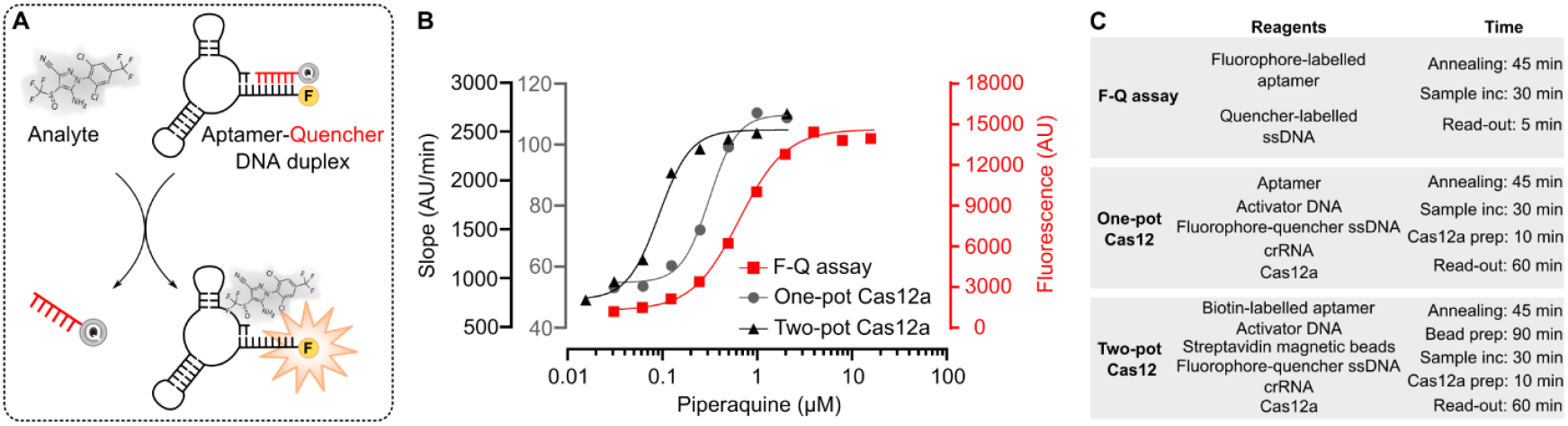
Aptasensor assay performance comparison. (A) Principle of the fluorophore-quencher displacement assay. (B) Comparison of the one-pot and two-pot CRISPR-Cas aptasensor assays (data are the same as shown in Figures 5E and 6C, respectively) with the F-Q assay. F-Q assay data represent two independent experiments each performed in duplicates; error bars are omitted for clarity. (C) Summary of reagent and time requirements for the three aptasensor assay formats.

This assay setup has been reported for two of the aptamer-analyte pairs included in this study, PQ4-piperaquine and FipA6B-fipronil.^30,31^ To obtain a direct comparison to the CRISPR-Cas aptasensor assay we thus aimed to reproduce these two F-Q assays exactly as previously reported.

For the FipA6B-fipronil aptamer-analyte pair, we did not observe any fluorescence signals upon incubating the aptamer-quencher duplex with fipronil. This was despite using the same DNA sequences labelled with the same fluorophore and quencher molecules, as well as following the exact F-Q assay procedure, as originally described.^30^

For the PQ4-piperaquine aptamer-analyte pair, we could closely reproduce the originally published F-Q assay data and obtained a sigmoidal dose-response curve over a range of 10 to 0.05 µM piperaquine (Figure 7B).^31^ LOD and LOQ were determined as 0.114 and 0.236 μM, respectively. We compared these data to both the one-pot and two-pot CRISPR-Cas aptasensor assay data for piperaquine. The two-pot assay could detect the lowest concentrations of piperaquine, followed by the one-pot assay and the F-Q assay. However, the differences were modest, with only two-to three-fold lower LOD and LOQ for the two-pot CRISPR-Cas aptasensor assay compared to the F-Q assay, which is less than what we expected given the enzymatic signal enhancement provided by the Cas12a assay. In Figure 7C, we show the reagents and approximate time requirements for the three assay setups, showing that the F-Q assay is the simplest, cheapest, and fastest to implement.

### Evaluation of aptamer-analyte interactions by isothermal titration calorimetry

Considering that for both MC-LR and fipronil we could not reproduce previously reported assay performances, we used isothermal titration calorimetry (ITC) to investigate the binding behavior between these analytes and their aptamers. As positive control we used the MN19-quinine aptamer-analyte pair, previously shown to elicit a strong signal in ITC.^34^ Indeed, we observed a strong ITC signal with MN19-quinine (Figure S7). In contrast, for both the AN6-MC-LR and FipA6B-fipronil aptamer-analyte pairs, we did not observe conclusive ITC signals that would indicate an enthalpically driven binding interaction. However, this alone is not decisive as to whether the two aptamers bind (or not) their analytes – complementary techniques such as nuclear magnetic resonance, mass spectrometry, or surface plasmon resonance would need to be employed to conclude on any interactions between these two aptamer-analyte pairs.^43^

## Discussion

Due to its simplicity, the CRISPR-Cas aptasensor assay is highly appealing because, in principle, it allows for fast, high-throughput sample analysis while only requiring a suitable aptamer-activator DNA pair and a fluorescence-capable photometer, e.g. in the form of a plate reader.

However, in our testing of this biosensor assay we came across several limitations, three of which we will discuss here in more detail: restrictions in the design of functional activator DNAs, performance versus workload of the CRISPR-Cas aptasensor assay, and problems with reproducibility of previously published aptamers and aptasensors.

First, the design of an activator DNA is a crucial step in the CRISPR-Cas aptasensor workflow. The aptamer-activator DNA duplex needs to be stable in absence of analyte but efficiently dissociate when the analyte is bound by the aptamer. A straightforward approach is to choose aptamers from the literature that were obtained by Capture-SELEX and then to redesign the capture strand as activator DNA. However, for the set of aptamers we chose to work with, we found that often the original capture strands had a relatively low affinity for the aptamer (that is, a low predicted melting temperature of the duplex) (Figure 2). This resulted in free (unbound) activator DNA, even when the activator DNA was annealed with a large molar excess of aptamer. A solution to this problem is to increase the number of complementary nucleotides between aptamer and activator DNA and hence raise the melting temperature of the duplex. This, however, may disrupt the delicate balance between aptamer-analyte binding and activator DNA release, to the effect that the activator DNA is bound too tightly to the aptamer and does not dissociate upon analyte binding.

A related complication is that optimal Cas12a activation requires an activator DNA with a length of 20 nt (Figure S4). However, Capture-SELEX strands are typically between 10 and 15 nt in length. We addressed this by adding “neutral” (non-annealing) nucleotides to the termini of the capture strands to arrive at 20 nt long activator DNAs (Figure 2). However, even if these nucleotides do not anneal to the aptamer, they may still influence the interaction between aptamer and activator DNA, or the aptamer and target analyte.

In our view, there is no straight-forward solution to these challenges. Essentially, activator DNA design remains a trial-and-error approach: each activator DNA design needs to be tested to determine whether it forms a sufficiently stable duplex with the aptamer while still releasing efficiently upon analyte binding. This becomes even more complicated if one wants to use an aptamer not obtained by Capture-SELEX, where the entire length of the aptamer needs to be scanned experimentally, ideally with a sliding window approach, to find a suitable site for activator DNA annealing and release. However, it is often not known *a priori* whether the aptamer is structure-switching upon analyte binding and whether activator DNA release will occur at all.

Considering this biochemically complex two-way interaction between aptamer-activator DNA and aptamer-analyte, detailed biochemical and structural information about each aptamer in consideration for the CRISPR-Cas aptasensor assay would be beneficial. However, such information about aptamers and their interaction with the analyte, including biochemical characterizations of the binding mode and structures of the aptamer in both the apo-form and ligand-bound state, are sparse and require detailed investigations.^43–45^ Potentially, more powerful 3D structure prediction and modelling tools may support a more rational approach to activator DNA design in the future.

A second point for discussion is the performance advantage, if any, of the CRISPR-Cas aptasensor. A purported benefit of this approach is the enzymatic signal amplification resulting in lower analyte limits of detection and quantification.^14,46^ Thus, we were surprised to find that, in our hands, the CRISPR-Cas aptasensor performed only moderately better than a simple fluorophore-quencher displacement assay, providing about two-to three-fold lower limits of analyte detection. At the same time, the CRISPR-Cas aptasensor assay comes with higher reagent costs and a more complicated and time-consuming workflow. Herein, we only have a direct comparison between the F-Q assay and the CRISPR-Cas aptasensor assays for one analyte (piperaquine), thus, it remains open whether for other aptamer-analyte pairs the difference in assay performance is more pronounced. In any case, the simple F-Q assay may be sufficient for many analyte detection purposes.

It has often been reported that CRISPR-Cas aptasensors provide a large linear detection range (Table S1). In the case of MC-LR, a span of six orders of magnitude was described^27^ - an astounding result given the underlying biochemical interactions, but one that we could not reproduce in this study. In our hands, the best performing aptasensor, targeting piperaquine, showed a linear range of about two orders of magnitude in both the one-pot and two-pot CRISPR-Cas aptasensor assays as well as in the F-Q assay. Such linear ranges of two to three orders of magnitude are typical and expected for bioassays based on a reversible, bimolecular binding reaction defined by K_d_, such as an aptamer-analyte interaction.^47^

Finally, a third point we would like to discuss are reproducibility problems with published aptamers and aptasensors. Our initial interest in the CRISPR-Cas aptasensor assay was sparked by a study targeting the cyanobacterial toxin MC-LR, which reported excellent assay performance and detection limits in the low femtomolar range.^27^ However, despite a significant number of attempts at assay improvement and testing, we did not succeed in observing any meaningful assay signal. Likewise, we could not reproduce a simple F-Q assay previously reported for the analyte fipronil.^30^ Both MC-LR and fipronil also did not generate a signal in ITC, although this does not unequivocally indicate lack of binding between aptamer and analyte since there are binding modes (*e*.*g*., primarily entropy-driven) that ITC is not sensitive to.

Of course, the fact that we failed to reproduce the MC-LR and fipronil assays does not imply that these assay do not, in principle, function as reported – it is possible that minute experimental details, which are not explicitly represented in the disclosed methods (*e*.*g*., differences in room temperature, minor impurities in buffers, different suppliers of reagents, etc.), determine whether the assays works or not. However, if this was the case, it would limit the robustness and thus general useability of such an assay. For the remaining non-functioning aptamer-analyte pairs tested (targeting ofloxacin, levofloxacin, and atrazine) we attempted to establish the CRISPR-Cas aptasensor setup *de novo*. The fact that these attempts did not work could be due to the limitations regarding activator DNA design, as discussed above. For example, for the LxCsh aptamer, shown clearly by ITC to bind its target levofloxacin,^33^ we had to use an activator DNA with a longer annealing region compared to the original Capture-SELEX strand in order to form a stable aptamer-activator DNA duplex. However, this likely also abolished activator DNA release upon target binding (or impacted target binding directly).

Reproducibility challenges and shortcomings of certain aptamers are well documented in the literature.^43,48–50^ This poses difficulties for developing new biosensors, as it is often unclear whether aptamers selected from the literature will function as intended. Considerable efforts for initial assay establishment and testing may therefore be necessary before realizing that a given aptamer does not perform reliably. Detailed binding studies at the start of an aptasensor project or generating entirely new aptamers oneself, for example by Capture-SELEX, are possible solutions but each represents a substantial experimental undertaking. The scarcity of functional and reliable aptamers is reflected in the “thrombin problem”, the observation that hundreds of aptasensors against the same few analytes (most prominently, thrombin) have been developed simply because the aptamers binding these analytes indeed function as expected and are thus used over and over again to showcase biosensor designs.^51^ Our results suggest that the PQ4 piperaquine aptamer,^31^ which functioned reproducibly well in our hands, could become another example of such a robust but niche aptamer upon which a large number of redundant aptasensors may be built in the future. At the same time, aptasensor assays for other relevant analytes are not being developed or fail to deliver reproducible results, calling into question the propagated potential of aptamers for biosensor applications.

In conclusion, we believe that the CRISPR-Cas aptasensor assay, on paper, is a great addition to the biosensor repertoire. In this study, however, this assay worked for only two out of seven analytes tested. Of course, we cannot rule out that much more extensive assay fine tuning for each single aptamer-analyte pair may have resulted in a higher success rate. However, many laboratories may not have the time and capacities required to engage in such additional assay fine-tuning, thereby limiting the useability and attractiveness of this approach compared to tried-and-proven analyte detection technologies such as mass spectrometry.

The Cas12a fluorescence assay itself worked flawlessly and could detect low picomolar concentrations of activator DNA, in line with its reported power in DNA detection.^52^ As we see it, the culprit of the CRISPR-Cas aptasensor workflow is the difficulty in finding reliable aptamers that indeed bind their intended analytes and efficiently release the activator DNA upon target binding. Furthermore, compared to a simple fluorophore-quencher displacement aptasensor, the added performance benefits of the CRISPR-Cas aptasensor were modest. Thus, the decision to implement a CRISPR-Cas aptasensor assay requires careful weighing of its added value against more established or simpler analyte detection systems.

## Materials and Methods

### Reagents and equipment

ssDNA oligonucleotides were purchased from IDT or Microsynth (see Table S2 for all oligonucleotides used in this work). crRNAs were purchased from IDT. All ssDNA or ssRNA oligonucleotides were dissolved in TE buffer (10 mM Tris, 0.1 mM EDTA, pH 7.5) and stored at −20°C. Buffer components were purchased from Sigma-Aldrich. MC-LR was purchased from Enzo Life Sciences (Cat.# ALX-350-012), atrazine was purchased from Sigma-Aldrich (Cat.# 45330), fipronil was purchased from Thermo Fisher Scientific (Cat.# 465570050), ofloxacin was purchased from Sigma-Aldrich (Cat.# O8757), levofloxacin was purchased from Thermo Fisher Scientific (Cat.# J66943.03), piperaquine was purchased from MedChemExpress (as tetraphosphate tetrahydrate; Cat.# HY-B1896B), and quinine was purchased from Thermo Fisher Scientific (as monohydrochloride dihydrate; Cat.# 163720050). Streptavidin magnetic beads (Cat.# S1420S) and Cas12a (Cat.# M0653T) were purchased from New England Biolabs. Oligonucleotide annealing was performed in a Biometra T3000 PCR machine, while temperature gradient incubations were performed in a Biometra T Professional Basic Gradient PCR machine. Fluorescence assays were read in a BioTek Synergy MX plate reader, controlled by Gen5 software. Melting temperatures of aptamer-activator DNA duplexes were calculated using the IDT OligoAnalyzer web tool.

### Cas12a fluorescence assay setup

Cas12a reaction setups for fluorescence read-out in a multiwell plate reader were prepared as follows: First, 9 μL of 10x CXB buffer were added to a well of a flat-bottom, clear 96-well plate. Next, 76 μL of sample to be analyzed (in water or buffer, depending on the experiment) were added per well. We then prepared 10 μL of Cas12a assay solution; this solution (or multiples thereof, depending on the number of samples to be analyzed) was prepared by mixing 7 μL ddH_2_O, 1 μL 10x CXB buffer, 1 μL crRNA (1 μM stock; matching the activator DNA in the sample to be analyzed), and 1 μL Cas12a (1 μM stock). This solution was incubated at 30°C for 3 min to ensure Cas12a-crRNA complex formation, and 10 μL were added per well. Lastly, we added 5 μL of fluorophore-quencher ssDNA reporter (10 μM stock) per well. Thus, each well contained, in a volume of 100 μL, a 1x concentration of buffer CXB (plus any buffer present in the 76 μL of sample solution), 10 nM crRNA-Cas12a complex, and 500 nM of fluorophore-quencher ssDNA reporter. The plate was tapped gently to mix all components and immediately transferred to the plate reader for analysis. The analysis program consisted of 5 s of initial shaking (slow speed setting), followed by a kinetic analysis of fluorescence intensity (without plate shaking): fluorescence measurements every 2 min over 60 min total, using the FAM channel (484/530 nm excitation/emission wavelengths, respectively), with temperature set to 25°C. Following the run, the data were exported from the Gen5 program to MS Excel as tabular data of fluorescence intensity over time.

### Analysis of Cas12a assay data

The plate reader data were transferred to GraphPad Prism (version 9.0.0) for analysis and plotting. We used standard linear regression to derive the slope of fluorescence increase over time (expressed as AU/min). We typically restricted the linear regression analysis to those data points within a time window of 10 to 30 min of the assay. This was done to ensure that we only analyzed data recorded after the assay had fully equilibrated and before linearity of the data may be compromised, e.g. by substrate limitations or enzyme inactivation. In some cases, e.g. with very high activator DNA concentrations during the LOD titrations, which reached the maximum fluorescence level detectable by the plate reader (100,000 AU) before 30 min, the linear regression analysis window was shifted accordingly. For the sigmoidal dose-response data obtained for the piperaquine F-Q assay and one-pot and two-pot CRISPR-Cas aptasensor data, we performed a 4PL nonlinear regression curve fit in Prism. LOD and LOQ were calculated by taking the resulting bottom asymptote value and adding 3 times or 10 times (for LOD and LOQ, respectively) the standard deviation of the matching blank samples. From the resulting y values, the corresponding x-values (*i.e*., analyte concentrations) were derived in Prism by interpolating from the curve fit.

### Determination of activator DNA LOD and LOQ

To determine the LOD and LOQ of each activator DNA we performed a dilution series from 4 nM to 1.95 pM (twelve 2-fold dilution steps). The dilution series was performed in 1x CXB buffer, followed by a standard Cas12a fluorescence read-out. For each activator DNA, the dilution series was repeated twice (experimental replicates), and for each dilution step two Cas12a reactions were performed (technical replicates) to yield a total of 4 datapoints per activator DNA concentration. The data were analyzed by linear regression, and we plotted the slope (AU/min) versus activator DNA concentration. The linear part (typically from 1.95 pM to 125 pM) of the resulting data was again analyzed by linear regression. From the resulting slope (s) and error (σ) of the regression line we calculated LOD and LOQ, defined as 3.3*σ/s and 10*σ/s, respectively.

### Annealing of activator and aptamer DNA

To anneal aptamers and activator DNAs we used a controlled temperature ramp program in a PCR machine to ensure reproducibility between experiments. Aptamer and activator DNA were mixed at the required concentrations and in the required buffer (both depending on the experiment) and distributed to PCR tubes. The tubes were transferred to the PCR machine and the annealing program was run, consisting of 3 min at 95°C, followed by a temperature ramp (0.03°C/s) to 25°C (approx. 40 min duration). The resulting aptamer-activator DNA complexes were used immediately thereafter; we did not store annealed complexes for several subsequent experiments, but annealing was performed freshly before each individual experiment.

### Titration of activator and aptamer DNA

To determine the efficiency of aptamer – activator DNA duplex formation we annealed a fixed concentration (10 nM) activator DNA with different concentrations of matching aptamer (0, 10, 20, 40, or 80 nM), using the standard annealing procedure and suitable buffer for each aptamer-activator DNA combination. Next, 76 μL of the resulting samples were directly transferred to the Cas12a fluorescence assay to gain insight into the relative amounts of remaining free activator DNA.

### Effect of target analytes on the Cas12a fluorescence assay

To test the effect of target analytes on the Cas12a fluorescence assay we set up standard Cas12a fluorescence assays with the R12.45 activator DNA (250 pM concentration). The assays were spiked with target analytes (or mock), typically at 50 to 100 μM concentration (or a 2-fold dilution series from 100 μM to 12.5 μM for piperaquine and quinine), followed by a standard fluorescence read-out. The data were visually analyzed for effects of the different target analytes on fluorescence generation.

### One-pot aptasensor Cas12a assay

For the one-pot aptasensor Cas12a assay, we annealed 25 nM activator DNA with 200 nM aptamer (1:8 molar ratio) in the matching buffer for each aptamer – analyte combination (Table S3); the total volume was chosen according to the number of samples to be analyzed. 250 μL aliquots were then distributed to PCR tubes and subjected to the annealing program, followed by re-pooling of all tubes. Next, 198 μL of the annealed aptamer-activator DNA solution were transferred to fresh PCR tubes and 2 μL of analyte stock solution was added; the analyte stock solution was prepared at a 100-fold concentration in order to reach the desired final analyte concentration, and co-solvent concentration (DMSO, methanol, acetone, buffer, or water, depending on the analyte) was 1% in the final sample. Negative control samples included only the co-solvent without analyte. The samples were then incubated at the desired temperature and the desired time, typically, between 20 and 50°C and for 30 min to 2h. Next, two times 76 μL from each tube were directly transferred to the Cas12a fluorescence assay for analysis.

### Two-pot magnetic bead aptasensor Cas12a assay

For the two-pot aptasensor Cas12a assay, we initially tried to reproduce the exact assay conditions and setup as reported for MC-LR by Kang and colleagues.^27^ When this was unsuccessful, we went through a stepwise assay modification procedure, including changes in buffers, reagent concentrations, and incubation times and temperatures, however, for MC-LR this remained unsuccessful. For piperaquine, the analyte for which this assay setup produced meaningful results, we settled on the following experimental procedure: 5 μL of streptavidin magnetic bead slurry (or multiples thereof), corresponding to 0.02 mg of beads and a manufacturer-stated 10 pmol biotinylated ssDNA binding capacity, were added to a PCR tube. The beads were washed 3x with 100 μL buffer AB, using a magnetic rack to separate the beads from solution, and finally resuspended in 50 μL AB. Note that, in case of multiple samples, we processed up to 10 times the amount of bead slurry (i.e., 50 μL) with the same procedure and wash volumes. In parallel, 150 nM aptamer and 150 nM activator DNA (or multiples thereof) were annealed in 200 μL buffer AB, corresponding to 30 pmol (or multiples thereof) of aptamer-activator DNA complex. Following the annealing procedure, the 200 μL of annealed complex were added to the 50 μL of washed beads, mixed by pipetting, and incubated for 30 min at room temperature. Next, the tube was placed on the magnetic rack, the supernatant was removed, and the beads were washed 5 times with 200 μL buffer AB. After the final wash the supernatant was removed; if functionalized beads for multiple samples were prepared, the beads were resuspended in 200 μL buffer AB and distributed accordingly (e.g., if beads for 10 samples were prepared, we distributed 10 times 20 μL of washed beads to fresh tubes). The supernatant was removed, and 100 μL of sample, containing analyte or mock controls, was added to the beads. The solution was mixed by pipetting and incubated for 30 min at room temperature. Finally, the tubes were placed on the magnetic rack, and 76 μL of supernatant were carefully removed and directly transferred to the Cas12a fluorescence assay.

### Fluorophore-quencher aptasensor assay

For the fluorophore-quencher displacement assay (F-Q assay), we followed the procedures as described for piperaquine and fipronil, respectively.^30,31^ For piperaquine, 100 nM FAM-labelled PQ4 aptamer was annealed with 500 nM Dabcyl-labelled quencher in PBS, 2 mM MgCl_2_, using our standard annealing protocol; the total volume was chosen according to the number of samples to be analyzed. Next, 40 μL of the annealed complex was mixed with 40 μL of piperaquine or mock sample (both in PBS, 2 mM MgCl_2_) in a well of a clear flat-bottom 96-well plate, and the plate was incubated for 30 min at room temperature in the dark. Finally, a single FAM fluorescence measurement was performed in the plate reader, the data were exported to MS Excel and analyzed in GraphPad Prism. For fipronil, 50 nM FAM-labelled FipA6B aptamer was annealed with 250 nM dabcyl-labelled quencher in 20 mM HEPES, 1 M NaCl, 10 mM MgCl_2_, 5 mM KCl, pH 7.5, using our standard annealing protocol. Next, 60 μL of the annealed complex was mixed with 60 μL of fipronil or mock samples (in the same buffer) in a well of a clear flat-bottom 96-well plate, and the plate was incubated for 40 min at room temperature in the dark. Finally, a single FAM fluorescence measurement was performed in the plate reader, the data were exported to MS Excel and analyzed in GraphPad Prism.

### Isothermal titration calorimetry

ITC was performed on a Malvern Panalytical Microcal iTC200 instrument and the matching software. Experiments were performed at 25°C, a stirring speed of 750 rpm, with an initial injection volume of 1.2 μL followed by 19 injections of 2 μL, and a spacing between injections of 90 to 180 s. Aptamers AN6 and FipA6B were used at 2 μM, while aptamer MN19 was used at 3.5 μM; aptamer solutions were placed in the sample cell. Matching analyte solutions (MC-LR, fipronil, or quinine, respectively) at 10-fold concentration of the aptamers were titrated into the aptamer solution or buffer; the matching buffer for each aptamer was used (Table S3), adjusted for co-solvent depending on the analyte. Data were analyzed with the iTC200 software.

## Data availability

All data are included in the manuscript and the supporting information.

## Author contributions

OFB, EMLJ and OTS designed the study; OFB performed experiments and analyzed data; OFB wrote the manuscript; OFB, EMLJ and OTS revised the manuscript; all authors read and approved the final manuscript.

## Acknowledgements

This work was supported by Eawag, the Swiss Federal Institute of Aquatic Science and Technology. Additionally, it was funded by the Swiss National Science Foundation’s Innosuisse funding scheme (103.792 IP-LS). We thank Dr. Ivan Corbeski at the Department of Biochemistry, University of Zurich, for support with ITC measurements. We also thank the Laboratory for Water Quality of the canton of Zurich (AWEL), the Department of the Environment of the canton of Aargau, the Laboratory for Environment and Energy of the canton of Lucerne, the Laboratory for Water and Energy of the canton of St. Gallen, and the Laboratory for Water and Soil Protection of the canton of Bern, for helpful discussions regarding biosensor implementation for cyanobacterial toxins.

