## Supplementary Information for "A critical assessment of aptamer and CRISPR-Cas12a-based biosensors for small molecule detection"

**Supplementary Information contents:**

Supplementary Tables p. 2

Supplementary Figures p. 8

Supplementary References p. 13

#### Supplementary Tables

#### Table S1

Selected CRISPR-Cas aptasensor studies published between 2019 and 2024.[a]

| Analyte | LOD | Linear Range | Assay Setup [b] | Ref |
|---|---|---|---|---|
| Kanamycin | 1 pM | 1 pM - 100 nM | Analyte-induced activator release, dsDNA generation by CHA, Cas12a activation, ssDNA-invertase cleaved off beads, sucrose>glucose conversion, glucose-meter read-out | 1 |
| TGF-β1 | 0.2 nM | 1 - 20 nM | Aptamer=activator, less free aptamer in presence of analyte, less Cas12a active, less ssDNA probe cleavage (inverse electrochemical detection) | 2 |
| Polychlorinated biphenyls | 128 fg/mL | 500 fg/mL - 50 ng/mL | Analyte-induced activator release, dsDNA generation by CHA, Cas12a activation, ssDNA F-Q probe cleavage | 3 |
| Prostate-specific antigen | 0.16 ng/mL | 0.31 - 5 ng/mL | Aptamer=activator, less free aptamer in presence of analyte, less Cas12a active, less ssDNA probe cleavage (inverse fluorescence detection) | 4 |
| Cocaine | 33 pM | 330 - 1.65 x $10^5$ pM | Split aptamer, analyte induces complex formation, TdT DNA extension, Cas12a activation by DNA extension, ssDNA F-Q probe cleavage | 5 |
| SARS-CoV2 nucleocapsid protein | 16.5 pg/mL | 50 pg – 100 ng/mL | Analyte-induced activator release, Cas12a activation, MB-polyA ssDNA reporter cleaved (electrochemical detection) | 6 |
| 8-oxoguanine | 32.3 pM | 0.1 - 1 nM | Aptamer-blocker complex, analyte induces blocker release, aptamer=activator activates Cas12a, ssDNA F-Q probe cleavage | 7 |
| α-fetoprotein | 10 ng/mL | 0.01 - 100 µg/mL | Analyte-induced activator release, Cas12a activation, ssDNA-invertase cleaved off beads, sucrose>glucose conversion, glucose-meter read-out | 8 |
| Aβ40 / Aβ42 (amyloid) | 0.1 pg/mL | 1-1000 pg/mL | Analyte-induced activator release, Cas12a activation, ssDNA F-Q probe cleavage | 9 |
| MC-LR | 3 pg/L | 1 x $10^{-5}$ - 10 µg/L | Analyte-induced activator release, Cas12a activation, ssDNA F-Q probe cleavage | 10 |
| ATP | 246 nM | 1.56 - 50 µM | Aptamer-activator complex on nanoneedles inserted into cells; needles with free activator retrieved, Cas12a activation, ssDNA F-Q probe cleavage | 11 |
| ATP | 0.85 µM | 5 - 500 µM | Aptamer=activator, less free aptamer in presence of analyte, less Cas12a active, less ssDNA probe cleavage (inverse LFA detection) | 12 |
| 17β-estradiol | 180 fM | $10^{-12}$ - $10^{-8}$ M | Split aptamer=activator, complex formation in presence of analyte, less Cas12a activation, less ssDNA reporter cleavage (inverse LFA detection) | 13 |
| ATP | 8.6 µM | 10 - 1000 µM | Analyte-induced activator release, Cas12a activation, ssDNA F-Q probe cleavage | 14 |
| IFN-γ | 100 fg/mL | 100 fg - 10 ng/mL | ELISA-like: biotin-ssDNA hybridized with aptamer=activator and bound to streptavidin plate; analyte releases aptamer; Cas12 activated by remaining aptamer on plate > ssDNA F-Q probe cleavage (inverse fluorescence signal) | 15 |
| Tobramycin | 1.5 pM | 5 - 30 pM | Analyte binding to aptamer induces SDA to create ssDNA activator, Cas12a activation, ssDNA F-Q probe cleavage | 16 |
| α -fetoprotein | 0.17 ng/mL | 0.5 - $10^4$ ng/mL | Analyte induces trigger DNA release from aptamer, hybridization chain reaction creates activator DNA, Cas12a activation, ssDNA F-Q probe cleavage | 17 |
| SARS-CoV2 Np | 0.077 ng/mL | 0.05 - 125 ng/mL | Aptamer=activator, less free aptamer in presence of analyte, less Cas12a active, less ssDNA probe cleavage (inverse fluorescence and electrochemical detection) | 18 |
| Adenosine | 15.7 nM | 0.5 - 100 µM | Analyte-induced activator release, Cas12a activation, ssDNA probe cleavage, fluorescence detection | 19 |
| $Cd^{2+}$ | 60 pM | 100 - 5000 pM | Aptamer without analyte triggers SDA, activator ssDNA generated, Cas12a activation, ssDNA probe cleavage, fluorescence detection; signal shutoff if analyte present | 20 |
| ATP | 1 nM | 0.1 - 750 µM | Analyte induces trigger DNA release from aptamer, hybridization chain reaction, Cas12a activation, ssDNA F-Q probe cleavage | 21 |

| | | | | |
|---|---|---|---|---|
| Aflatoxin B1 | 0.004 ng/mL | 0.01 - 20 ng/mL | Analyte-induced activator release, Cas12a activation, ssDNA F-Q probe cleavage | 22 |
| Ochratoxin A | 1.56 ng/mL | 5 - 80 ng/mL | Analyte-induced activator release, Cas12a activation, ssDNA F-Q probe cleavage | 23 |
| ATP | 104 nM | 25 - 500 µM | Aptamer=activator, less free aptamer in presence of analyte, less Cas12a active, less ssDNA probe cleavage (inverse fluorescence detection) | 24 |
| ATP | 400 nM | 1 - 200 µM | Aptamer=activator, less free aptamer in presence of analyte, less Cas12a active, less ssDNA probe cleavage (inverse fluorescence detection) | 25 |
| Melamine | 38 nM | 0.1 - 25 µM | Analyte-induced activator release, Cas12a activation, ssDNA F-Q probe cleavage | 26 |
| ATP | 490 nM | 1 µM - 1 mM | ATP aptamer coupled to Cas12a inhibitory aptamer, analyte triggers conformational change to release inhibitory aptamer, separate dsDNA activates Cas12a, ssDNA F-Q probe cleavage | 27 |
| ATP | 0.2 µM | n.r. | Aptamer-activator sandwich, analyte releases activator, Cas12a activation, ssDNA-HRP cleaved off beads, HRP read-out | 28 |
| Exosomes (CD63 / EpCAM) | n.r. | $10^3$ - $10^7$<br>$10^4$ - $10^8$<br>particles/mL | Analyte-induced activator release, Cas12a/Cas13 activation, ssDNA F-Q probe cleavage, dual FAM/Cy5 fluorescence output | 29 |
| Aflatoxin B1 | 5.2 pg/mL | 50 pg/mL - 100 ng/mL | Analyte-induced activator release, Cas12a activation, ssDNA-HRP cleaved from within pipet tip, HRP read-out | 30 |
| Methamphetamine, Cocaine | 27.5 / 10.7 pg/mL | 0.1-100 ng/mL | Analyte-induced activator release, Cas12a activation, ssDNA F-Q probe cleavage | 31 |
| Estradiol E2, PSA | 0.04 / 0.65 pg/mL | 0.05 - 80<br>1 - 200 pg/mL | Analyte-induced activator release, activator triggers CHA, generated dsDNA activates Cas12a, ssDNA probe cleavage, fluorescence detection | 32 |
| B-type natriuretic peptide | 13.5 fM | 5 - 100 fM | Analyte-induced activator release, activator triggers entropy-driven catalysis, Cas12a activation, ssDNA probe cleaved (electrochemical detection) | 33 |
| MC-LR | 19 pg/mL | 50 pg - 1 µg/mL | Aptamer incubated with analyte, activator added, free activator activates Cas14, ssDNA-fluorophore cleaved from MOF quencher (fluorescence detection) | 34 |
| MC-LR | 4.5 pg/mL | 0.01 - 50 ng/mL | Aptamer incubated with analyte and activator, free activator activates Cas12a > ssDNA-HRP cleaved off MBs, HRP read-out | 35 |
| Aflatoxin B1 | 0.92 pg/mL | 0.001 - 80 ng/mL | Aptamer-activator sandwich, analyte-induced activator release, Cas12a activation, ssDNA-fluorophore cleaved off MXene quencher (fluorescence detection) | 36 |
| SARS-CoV2 spike protein 1 | 1.5 pg/mL | 5 pg/mL - 200 ng/mL | Aptamer=activator, less free aptamer in presence of analyte, less Cas12a active, less ssDNA probe cleavage (inverse electrochemical detection) | 37 |
| ATP | 0.21 uM | 0.78 - 25 µM | Aptamer-activator sandwich, analyte-induced activator release, Cas12a activation, ssDNA F-Q reporter cleavage | 38 |
| Ochratoxin A | 0.29 pg/mL | 1 - 5000 pg/mL | Aptamer=activator, less free aptamer in presence of analyte, less Cas12a active, less ssDNA probe cleavage (inverse electrochemical detection) | 39 |
| Acetamiprid/Atrazine | 2.5 / 0.2 pM | 10 pM - 1 uM<br>1 pM - 100 nM | Analyte-induced blocker DNA release from aptamer, SDA on aptamer generates activator dsDNA, Cas12a activation, ssDNA F-Q probe cleavage | 40 |
| Zearalenone | 0.21 pg/mL | 1 - 1000 pg/mL | Analyte-induced activator release, dsDNA formation with ssDNA 2, Nt.AlwI enzyme creates activator ssDNA 3, Cas12a activation, ssDNA F-Q probe cleavage | 41 |
| Ampicillin | 10 pM | 0.01 - 500 nM | Analyte-induced activator release, Cas12a activation, ssDNA F-Q probe cleavage | 42 |
| SARS-CoV2 S1 / ATP | 0.06 pM<br>0.06 µM | 0.2 pM - 10 nM<br>0.2 - 1000 µM | Aptamer with locker region forms hairpin, T4 polymerase elongation in absence of analyte creates dsDNA for Cas12a activation. With analyte: no hairpin formation, signal-off | 43 |
| Ochratoxin A | 38 fg/mL | 100 fg - 50 ng/mL | Analyte-induced activator release, Cas12a activation, ssDNA reporter cleavage (electrochemical detection) | 44 |
| 17β-estradiol, bisphenol A | 0.08 nM / 0.06 nM | 0.2 - 25 nM<br>0.1 - 250 nM | Aptamer-activator sandwich, analyte-induced activator release, Cas12a activation, ssDNA F-Q probe cleavage | 45 |

| | | | | |
|---|---|---|---|---|
| SARS-CoV2 NP | 32 fM | 0.19 - 781 pM | Analyte-induced activator release, Cas12a activation, ssDNA F-Q probe cleavage | [46] |
| ATP, $Cd^{2+}$, histamine, aflatoxin B1, thrombin | 80 nM<br>4 nM<br>30 nM<br>6 nM<br>36 nM | 100 - 2000 nM<br>5 - 1500 nM<br>40 - 1500 nM<br>20 - 3000 nM<br>40 - 2000 nM | Aptamer=activator, less free aptamer in presence of analyte, less Cas12a active, less ssDNA probe cleavage (inverse fluorescence detection) | [47] |
| Aflatoxin B1, $Cd^{2+}$ | 31 pM<br>3.9 nM | 0.25 - 31.3 nM<br>3.9 nM - 8 µM | Capture DNA on plate, hybridizes with aptamer=activator without analyte, Cas12a activation, ssDNA F-Q reporter cleavage. Signal-off in presence of analyte. | [48] |
| Thrombin | 10 pM | 10 - 625 pM | ELISA-like; analyte capture on plate by immobilized aptamer or antibody, secondary aptamer=activator binds, Cas12a activation, ssDNA F-Q reporter cleavage | [49] |

[a] In total, as of 2025, approximately 150 to 200 studies (depending on the exact search criteria) that report a combination of aptamers and CRISPR-Cas enzymes for biosensor design have been published since 2019.

[b] We provide here a short summary of often highly complex assay setups; please check the original publications for accurate assay descriptions.

Abbreviations: CHA, catalytic hairpin assembly; SDA, strand displacement amplification; HRP, horseradish peroxidase; MB, magnetic beads; MOF, metal-organic framework.

Aptamer=activator refers to setups where a part of the aptamer sequence is also the activator sequence (complementary to the crRNA) of Cas12a.

Aptamer-activator sandwich refers to setups where aptamer and activator are specifically designed to anneal in defined stoichiometries and geometries, typically two aptamers sequestering one activator DNA.

#### Table S2

DNA and RNA sequences used in this study.

| Name | Sequence 5' - 3' [a] | Ref |
|---|---|---|
| MC-LR aptamer AN6 | GGCGCCAAACAGGACCACCATGACAATTACCCATACCACCTCATTATGCCCCATCTCCGC | [53] |
| AN6 5'-Biotin | Bio-TEG-GGCGCCAAACAGGACCACCATGACAATTACCCATACCACCTCATTATGCCCCATCTCCGC | [53] |
| AN6 activator DNA 1 | CGAGCGGAGATGGGGCATGA | [53] |
| AN6 activator DNA 2 | GATGGGGCATAATGAGGTGG | [35] |
| AN6 crRNA 1 | UAAUUUCUACUAAGUGUAGAU<u>UCAUGCCCCAUCUCCGCUCG</u> | [53] |

| | | |
|---|---|---|
| AN6 crRNA 2 | UAAUCUACUAAGUGUAGAUCCACCUCAUUAUGCCCCAUC | [35] |
| Quinine aptamer MN19 | GACAAGGAAAATCCTTCAACGAAGTGGGTC | [55] |
| MN19 5'-Biotin | Bio-TEG-GACAAGGAAAATCCTTCAACGAAGTGGGTC | This work |
| MN19 activator DNA | CCACTTCGTTGAAGGATTTG | [57] |
| MN19 crRNA | UAAUCUACUAAGUGUAGAUCAAAUCCUUCAACGAAGUGG | [57] |
| Piperaquine aptamer PQ4 | CTCTCGGGACGACCACACGCTAGTTGCGCCGAATGGCCGTGAAAGTCGTCCC | [51] |
| PQ4 3'-Biotin | CTCTCGGGACGACCACACGCTAGTTGCGCCGAATGGCCGTGAAAGTCGTCCC-TEG-Bio | This work |
| PQ4 activator DNA | GGTCGTCCCGAGAGCAGTAG | This work |
| PQ4 crRNA | UAAUCUACUAAGUGUAGAUCUACUGCUCUCGGGACGACC | This work |
| 5'-FAM PQ4 aptamer | FAM-CTCTCGGGACGACCACACGCTAGTTGCGCCGAATGGCCGTGAAAGTCGTCCC | [57] |
| PQ4-dabcyl Quencher | GGTCGTCCCGAGAG-DABC | [57] |
| Ofloxacin aptamer Q2 | ATACCAGCTTATTCAATTGCAGGGTATCTGAGGCTTGATCTACTAAATGTCGTGGGGCATTGCTATTGGCGTTGATACGTACAATCGTAATCAGTTAG | [54] |
| Ofloxacin aptamer Q8 | ATACCAGCTTATTCAATTAGTTGTGTATTGAGGTTTGATCTAGGCATAGTCAACAGAGCACGATCGATCTGGCTTGTTCTACAATCGTAATCAGTTAG | [54] |
| Q2 Activator DNA | GTAGATCAAGCCTCACCGAC | This work |
| Q8 Activator DNA | CTAGATCAAACCTCACCGAC | This work |
| Q2/Q8 crRNA | UAAUUUCUACUAAGUGUAGAUGUCGGUGAGGYUUGAUCUAS | This work |
| Levoflo-xacin aptamer LxCsh | GGGUUUGGGUAGUGCGAUUCGCACUGAAUGCCGCGUAGGCUUCGCUACUGGCUUCUACUACGUACCCAAACCC | [56] |
| LxCsh activator DNA | GCGTAGAAGCCAGTAGCATC | This work |
| LxCsh crRNA | UAAUUUCUACUAAGUGUAGAUGAUGCUACUGGCUUCUACGC | This work |
| Fipronil aptamer FipA6B | CTCTCGGGACGACAGATAGTGTGTACATGAAGGGTCGTCCC | [50] |

| | | |
|---|---|---|
| Fipronil aptamer FipA8 | CTCTCGGGACGACACGGGGCGCCACTAATGATGTATTGGTGCGACAGGCGTCGTCCC | [50] |
| FipA8 activator DNA 1 | GTGTCGTCCCGAGAGCCGAC | This work |
| FipA6B activator DNA 1 | CTGTCGTCCCGAGAGCCGAC | This work |
| FipA8 / FipA6B activator DNA 2 | GTCGTCCCGAGAGCCGAGCT | This work |
| FipA8 / FipA6B crRNA 1 | UAAUUUCUACUAAGUGUAGAUGUCGGCUCUCGGGACGACAS | This work |
| FipA8 / FipA6B crRNA 2 | UAAUUUCUACUAAGUGUAGAUAGCUCGGCUCUCGGGACGAC | This work |
| FipA6B-FAM | FAM-CTCTCGGGACGACAGATAGTGTGTACATGAAGGGTCGTCCC | [50] |
| FipA6B-dabcyl quencher | GTCGTCCCGAGAG-DABC | [50] |
| Atrazine aptamer R12.45 | TGTACCGTCTGAGCGATTCGTACTTTATTCGGGAAGGGTATCAGCGGGG | [52] |
| R12.45 aptamer +1 | GTGTACCGTCTGAGCGATTCGTACTTTATTCGGGAAGGGTATCAGCGGGG | This work |
| R12.45 aptamer +3 | TAGTGTACCGTCTGAGCGATTCGTACTTTATTCGGGAAGGGTATCAGCGGGG | This work |
| R12.45 aptamer +5 | AATAGTGTACCGTCTGAGCGATTCGTACTTTATTCGGGAAGGGTATCAGCGGGG | This work |
| R12.45 aptamer +7 | CTAATAGTGTACCGTCTGAGCGATTCGTACTTTATTCGGGAAGGGTATCAGCGGGG | This work |
| R12.45 activator DNA | CAGACGGTACACTATTAGAC | This work |
| R12.45 activator 1-15 | GGTACACTATTAGAC | This work |
| R12.45 activator 4-18 | GACGGTACACTATTA | This work |
| R12.45 activator 7-20 | CAGACGGTACACTA | This work |
| R12.45 crRNA | UAAUUUCUACUAAGUGUAGAUGUCUAAUAGUGUACCGUCUG | This work |

| | | |
|---|---|---|
| Cas12a reporter 5 nt | FAM-TTATT-BHQ1 | [58] |
| Cas12a reporter 12 nt | FAM-TTTTTTTTTTTT-BHQ1 | [59] |

[a] The underlined part of the crRNA sequences is the region complementary to the matching activator DNA.

#### Table S3

Buffers used in this study.

| Buffer | Composition | Ref |
|---|---|---|
| Fipronil buffer | 20 mM HEPES, 1 M NaCl, 10 mM $MgCl_2$, 5 mM KCl, pH 7.5 | [50] |
| Piperaquine buffer | PBS, 2 mM $MgCl_2$ | [51] |
| Atrazine buffer | 20 mM Tris, 100 mM NaCl, 5 mM $MgCl_2$, pH 7.4 | [52] |
| MC-LR buffer | 50 mM Tris, 150 mM NaCl, 2 mM $MgCl_2$, pH 7.5 | [53] |
| Ofloxacin buffer | 20 mM Tris, 100 mM NaCl, 2 mM $MgCl_2$, 1 mM $CaCl_2$, pH 7.6 | [54] |
| Quinine buffer | 20 mM Tris, 140 mM NaCl, 5 mM KCl, pH 7.4 | [55] |
| Levofloxacin buffer | 40 mM HEPES, 20 mM NaCl, 5 mM $MgCl_2$, 0.25 mM KCl, pH 7.4 | [56] |
| TE buffer | 10 mM Tris, 1 mM EDTA, pH 8.0 | n.a. |
| PBS | 137 mM NaCl, 2.7 mM KCl, 10 mM $Na_2HPO_4$, 1.8 mM $KH_2PO_4$, pH 7.4 | n.a. |
| Annealing buffer | 10 mM Tris, 50 mM NaCl, 2 mM $MgCl_2$, pH 7.5 | This work |
| NEB2.1 | 10 mM Tris-HCl, 50 mM NaCl, 10 mM $MgCl_2$, 0.1 mg/ml recombinant albumin, pH 7.9 | NEB |
| CX buffer | 50 mM potassium acetate, 20 mM Tris acetate, 20 mM $MgSO_4$, 10 mM DTT, pH 8.0 | This work |
| CXB buffer | 50 mM potassium acetate, 20 mM Tris acetate, 20 mM $MgSO_4$, 10 mM DTT, 0.1 mg/mL BSA, pH 8.0 | This work |
| CM buffer | 50 mM potassium acetate, 20 mM Tris acetate, 10 mM $MnSO_4$, 10 mM DTT, pH 8.0 | This work |
| CMX buffer | 50 mM potassium acetate, 20 mM Tris acetate, 10 mM $MnSO_4$, 10 mM DTT, 0.1 mg/mL BSA, pH 8.0 | This work |

### Supplementary Figures

#### Figure S1

```
PQ4          5'-UAAUUUCUACUAAGUGUAGAUcuacugcucucgggacgacc-3'
                                     ||||||||||||||||||||
                                  3'-GATGACGAGAGCCCTGCTGG-5'

MN19         5'-UAAUUUCUACUAAGUGUAGAUcaaauccuucaacgaagugg-3'
                                     ||||||||||||||||||||
                                  3'-GTTTAGGAAGTTGCTTCACC-5'

R12.45       5'-UAAUUUCUACUAAGUGUAGAUgucuaauaguguaccgucug-3'
                                     ||||||||||||||||||||
                                  3'-CAGATTATCACATGGCAGAC-5'

FipA8        5'-UAAUUUCUACUAAGUGUAGAUagcucggcucucgggacgac-3'
FipA6B                               ||||||||||||||||||||
                                  3'-TCGAGCCGAGAGCCCTGCTG-5'

AN6 Act.1    5'-UAAUUUCUACUAAGUGUAGAUucaugccccaucuccgcucg-3'
                                     ||||||||||||||||||||
                                  3'-AGTACGGGGTAGAGGCGAGC-5'

AN6 Act.2    5'-UAAUUUCUACUAAGUGUAGAUccaccucauuaugccccauc-3'
                                     ||||||||||||||||||||
                                  3'-GGTGGAGTAATACGGGGTAG-5'

Q2           5'-UAAUUUCUACUAAGUGUAGAUgucggugaggyuugaucuas-3'
Q8                                   ||||||||||||||||||||
                                  3'-CAGCCACTCCGAACTAGATG-5'
                                  3'-CAGCCACTCCAAACTAGATC-5'

LxCsh        5'-UAAUUUCUACUAAGUGUAGAUgauccuacuggcuucuaggc-3'
                                     ||||||||||||||||||||
                                  3'-CTAGGATGACCGAAGATCCG-5'
```

**Figure S1. crRNAs used in this study**. The nucleotide sequences of crRNAs used in this study are shown together with the sequences of their corresponding activator DNAs (red).

#### Figure S2

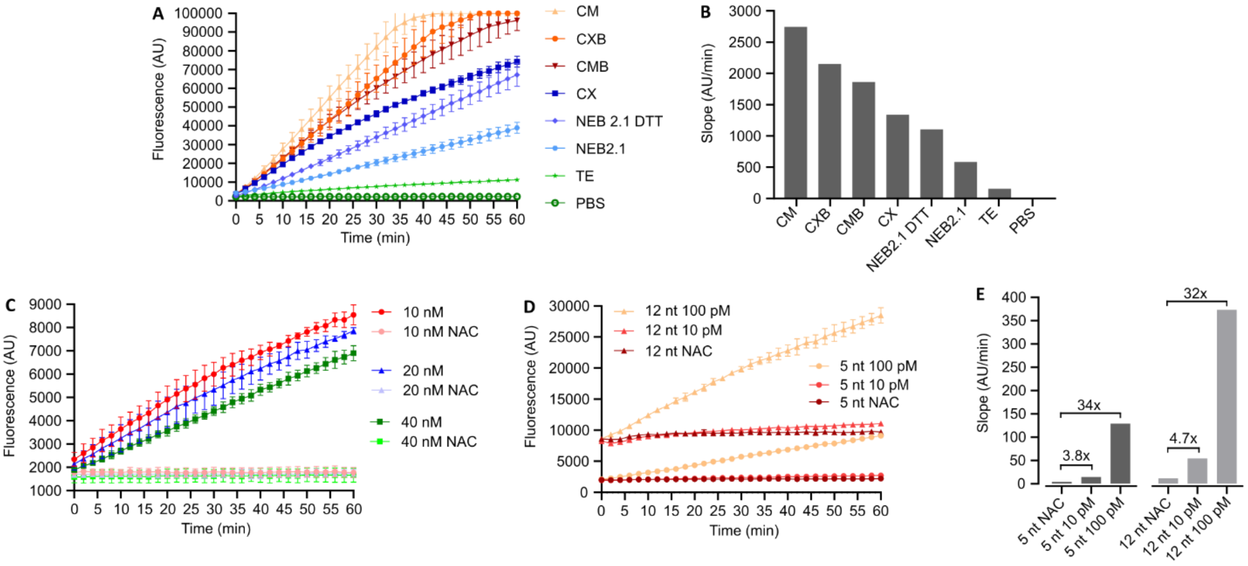


**Figure S2. Cas12a fluorescence assay optimization.** (A) Comparison of the Cas12a fluorescence assay set up with different buffers. Assays were performed with 1 nM activator DNA. (B) Slope of fluorescence increase over time derived from the data shown in (A). (C) Comparison of fluorescence generation by different amounts of Cas12a-crRNA complexes. Assays were performed with 100 pM activator DNA; NAC, no activator DNA control. (D) Comparison of fluorescence generation by Cas12a using a 12-nt- or 5-nt-long fluorophore-quencher ssDNA reporter. Assays were performed with 100 pM or 10 pM activator DNA. (E) Slope of fluorescence increase over time derived from the data shown in (D); the fold-change in slope over the negative controls is indicated.

#### Figure S3

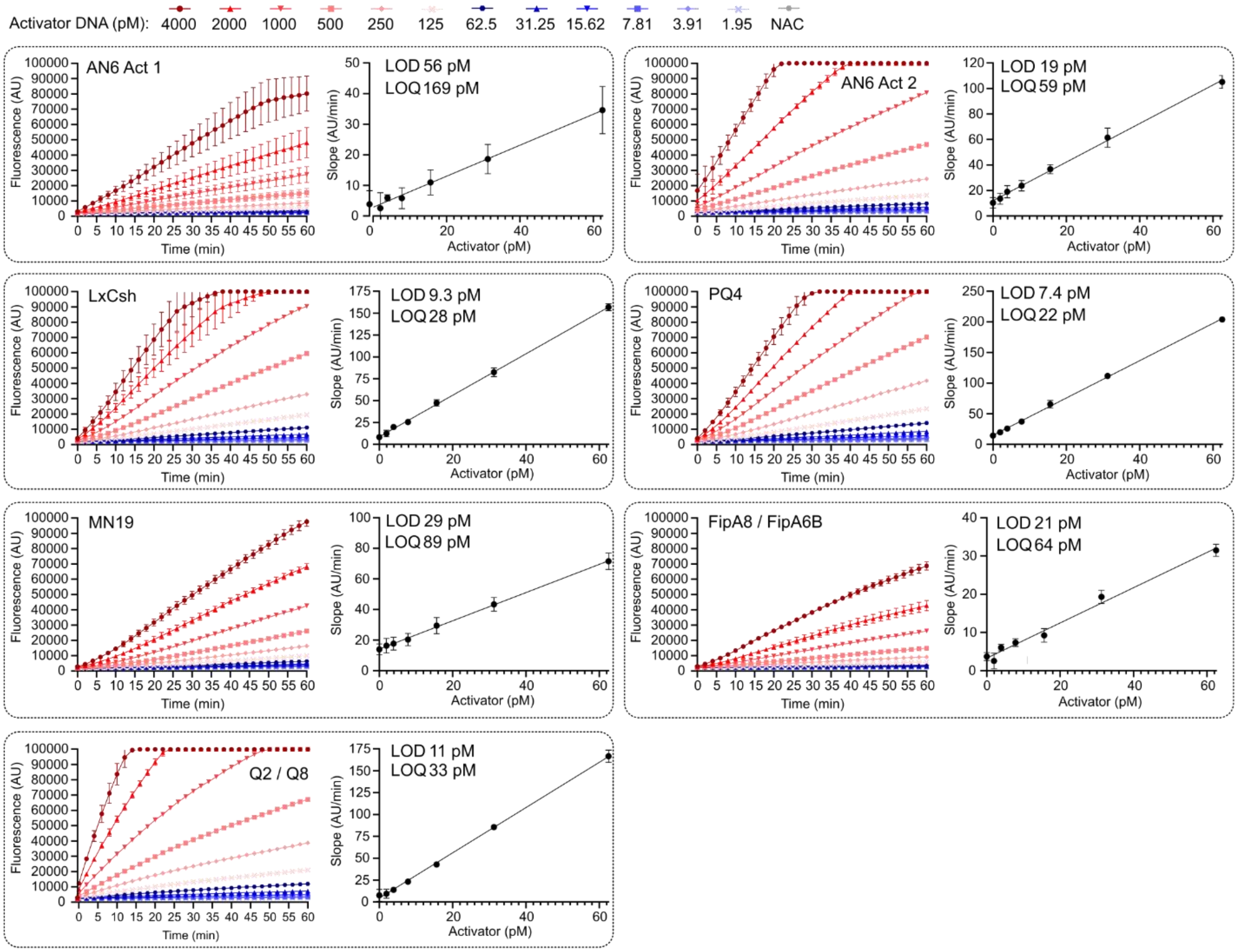


**Figure S3. LOD and LOQ determination for all activator DNAs.** For each activator DNA the original fluorescence data from the Cas12a assay (derived from two independent experiments, each performed in duplicates) and the resulting linear regression line used to determine the LOD and LOQ values, are shown.

#### Figure S4

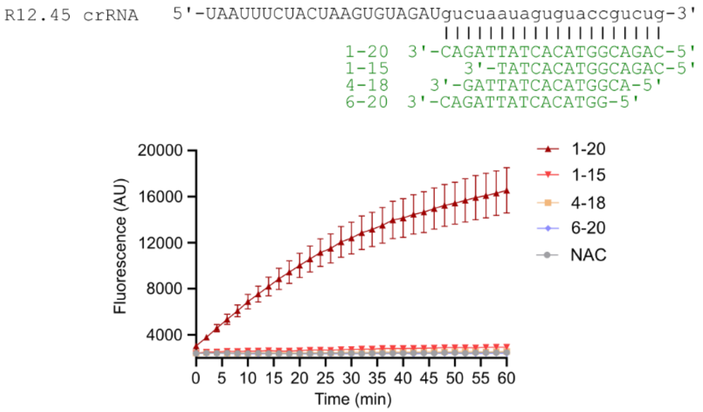


**Figure S4. Activator DNA length requirements.** Several shorter versions of the original 20 nt atrazine activator DNA were compared for their Cas12a activation potential, revealing that they all fall short compared to the full-length activator DNA.

#### Figure S5

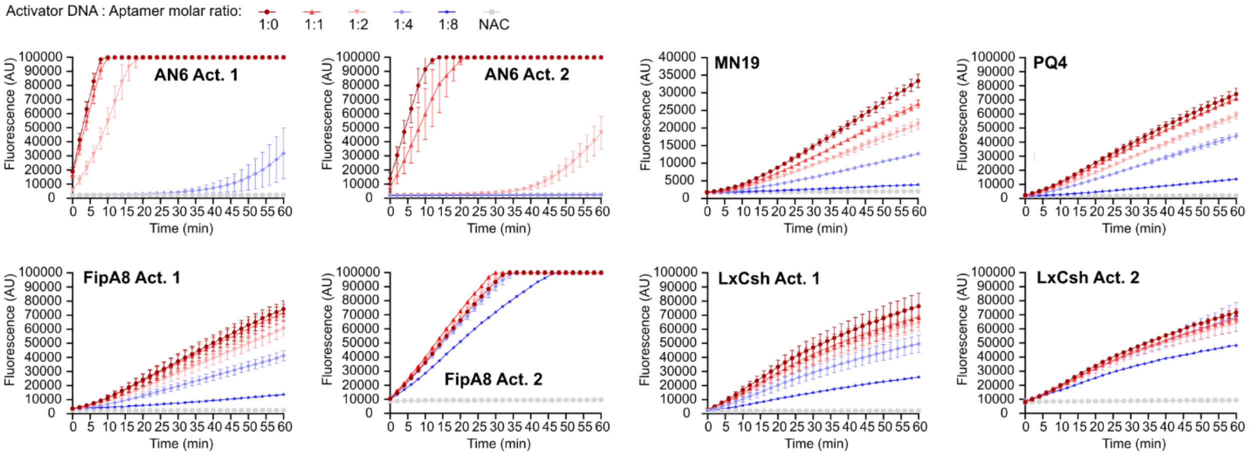


**Figure S5. Titration series of all aptamer-activator DNA pairs.** Shown are titration series of varying molar ratios of aptamer and the corresponding activator DNA for all aptamer-activator DNA pairs, aimed at determining the conditions under which minimal free activator DNA remains in solution, as measured by the Cas12a fluorescence assay. For FipA8/FipA6B and LxCsh, we also tested activator DNAs (Act. 2) designed based on the original Capture-SELEX capture strands; however, they failed to form stable duplexes with the aptamer, even at 8-fold molar excess of aptamer.

#### Figure S6

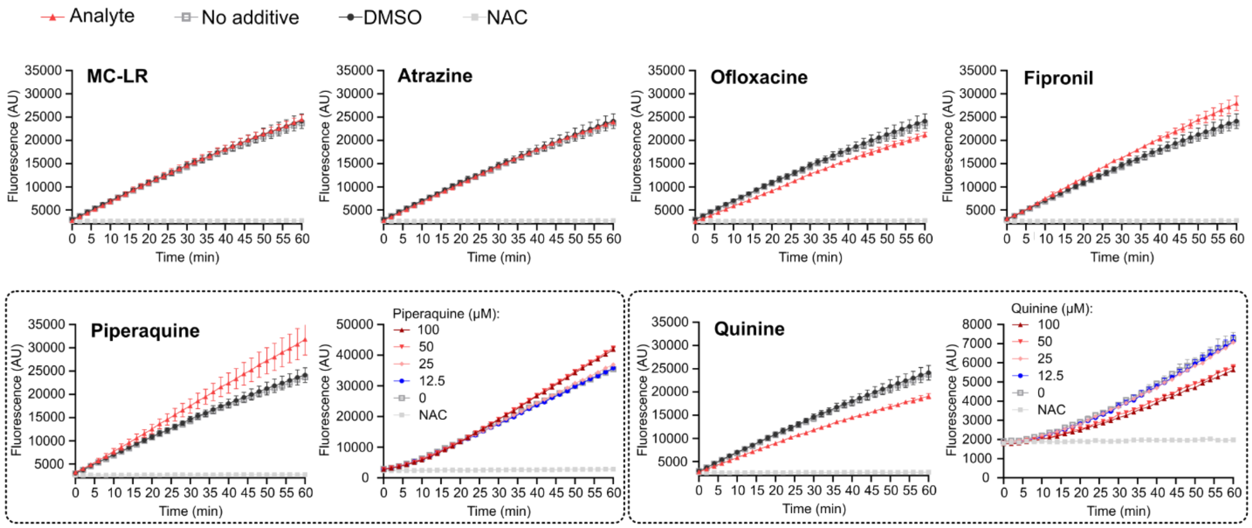


**Figure S6. Effect of analytes on the Cas12a fluorescence assay.** The effect of each analyte on fluorescence generation in the Cas12a assay was tested (MC-LR: 50 µM; all other analytes 100 µM). For piperaquine and quinine, we observed a mild positive and negative effect on fluorescence generation, respectively. Dilution series indicated that for both substances, this effect diminished below concentrations of 25 µM. Levofloxacin was not tested, as this stereoisomer is included in the racemic mixture of ofloxacin. The Cas12a assay was set up using the R12.45 activator DNA.

#### Figure S7

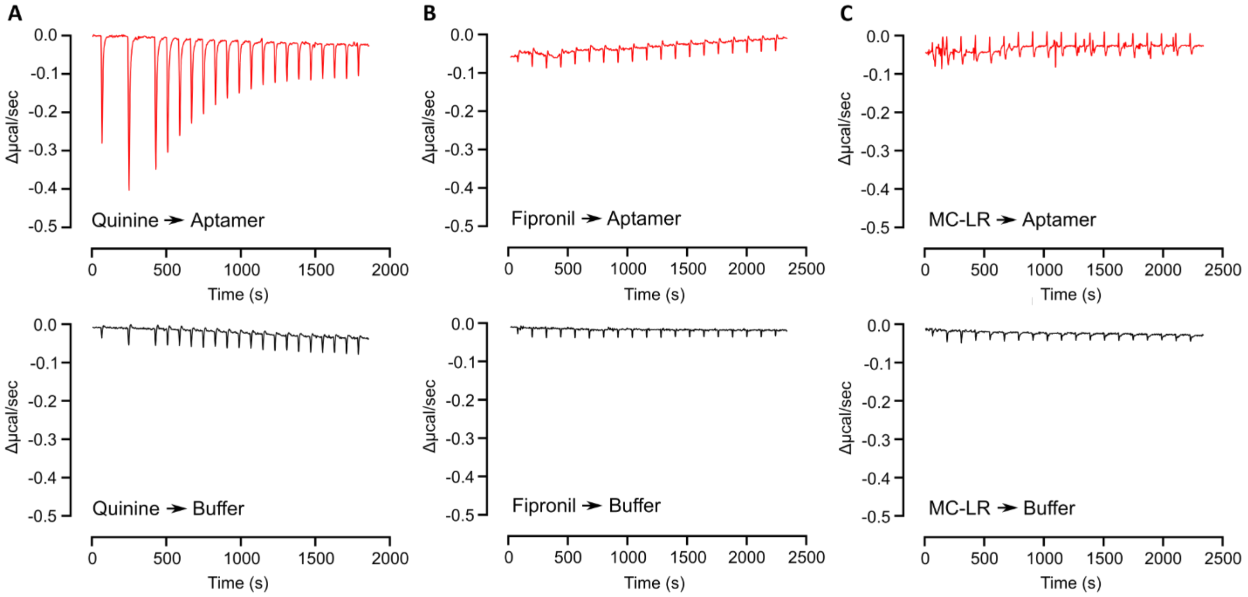


**Figure S7. Isothermal titration calorimetry to investigate aptamer-analyte binding.** ITC was performed with the quinine-MN19 (A), fipronil-FipA6B (B), and MC-LR-AN6 (C) analyte-aptamer pairs. Shown are the original ITC data obtained by titrating the analyte into aptamer solution (top row) and the matching titration of analyte into buffer to obtain the heat of dilution (bottom row). The data were scaled to show only the change in µcal/sec for better comparability. Data from one exemplary titration are shown per analyte; the titrations were repeated two to three times with comparable results.
